# Synaptic wiring motifs in posterior parietal cortex support decision-making

**DOI:** 10.1101/2022.04.13.488176

**Authors:** Aaron T. Kuan, Giulio Bondanelli, Laura N. Driscoll, Julie Han, Minsu Kim, David G. C. Hildebrand, Brett J. Graham, Logan A. Thomas, Stefano Panzeri, Christopher D. Harvey, Wei-Chung A. Lee

## Abstract

The posterior parietal cortex (PPC) exhibits choice-selective activity during perceptual decision-making tasks. However, it is not known how this selective activity arises from the underlying synaptic connectivity. Here, we combined virtual reality behavior, two-photon calcium imaging, high throughput electron microscopy, and circuit modeling to analyze how synaptic connectivity between neurons in PPC relates to their selective activity. We found that excitatory pyramidal neurons preferentially target inhibitory interneurons with the same selectivity. In turn, inhibitory interneurons preferentially target pyramidal neurons with opposite selectivity, forming an opponent inhibition motif. Using circuit models, we show that opponent inhibition amplifies selective inputs and induces competition between neural populations with opposite selectivity, thereby improving the encoding of trial-type information. These results provide evidence for how synaptic connectivity in cortical circuits supports a learned decision-making task.

## Introduction

Decision-making is a critical component of behavior and cognition, and understanding how it is implemented has been a long-standing goal in neuroscience. Experiments with primates have revealed the importance of the neocortex for perceptual decision-making, including the posterior parietal cortex (PPC), where neuronal activity is predictive of upcoming behavioral choices^1–3^. Such choice-selective activity has also been found in rodent PPC^4–12^, but it remains unclear how this selective neuronal activity arises and to what extent it is generated from local synaptic connectivity.

Models of cortical decision-making circuits propose that choice alternatives are represented by pools of recurrently connected excitatory neurons, which compete via inhibitory connectivity^13,14^. In some cases, inhibitory connectivity is assumed to be non-selective and random, but recent work has shown that inhibitory activity is as selective as excitatory activity, which suggests that inhibitory connectivity may follow choice-selective rules^10^. Indeed, recent models have shown that selective inhibition can crucially alter circuit function by stabilizing network activity or maximizing competition between opposing excitatory pools^15^. However, activity measurements in PPC are consistent with multiple possible circuit architectures^10,15,16^, and the underlying synaptic connectivity has not previously been measured.

Until recently, direct measurements of synaptic connectivity within large neuronal populations have not been technically feasible. However, advances in high-throughput electron microscopy (EM) have now made it possible to comprehensively map synaptic connectivity within circuits^17–25^. Such connectomic approaches in cortex have focused mainly on sensory areas such as visual cortex^26–30^, where interneuron activity and connectivity is generally less selective than excitatory^23,31^ (but see ^32–34^). As a result, little is known about synaptic connectivity in association areas such as PPC or how it may differ from sensory cortex.

Here, we combined a decision task, 2-photon calcium imaging, and automated serial-section EM^4,17,35^ to measure how synaptic connectivity of hundreds of cortical neurons relates to their functional selectivity in PPC. We found selective excitatory-to-inhibitory (E-to-I) and inhibitory-to-excitatory connectivity (I-to-E): excitatory neurons preferentially targeted inhibitory neurons with the same selectivity, while inhibitory neurons preferred excitatory targets with opposite selectivity. Together, these preferences form an opponent inhibition motif, in which the activity of left-selective excitatory neurons suppresses the activity of right-selective ones, and vice versa. To investigate the functional implications of this connectivity motif, we modeled recurrent circuits with excitatory and inhibitory populations and found that opponent inhibition supports competition between opposing pools and amplification of input selectivity, which promotes reliable encoding of the trial type.

## Results

### Behavior, Functional Imaging, and EM

We trained mice to perform a two-alternative forced-choice task in a virtual reality T-maze and used 2-photon calcium imaging to measure activity of layer 2/3 neurons in left-hemisphere PPC during task performance^4,35^ (Fig. 1a, Ext. Data Fig. 1a-c). These behavioral and functional imaging data were included in a previous study^35^. Consistent with previous results^4,10,35^, we found that many PPC neurons exhibited activity that was selective for trial-type (left or right turn trials, Fig. 1b, Supp. Fig. 1). To quantify this selectivity, we defined a selectivity index (SI) based on the peak mutual information between the neuronal activity and trial-type across the maze (Fig. 1b, Ext Data Fig. 1d, Methods).

**Fig. 1:**
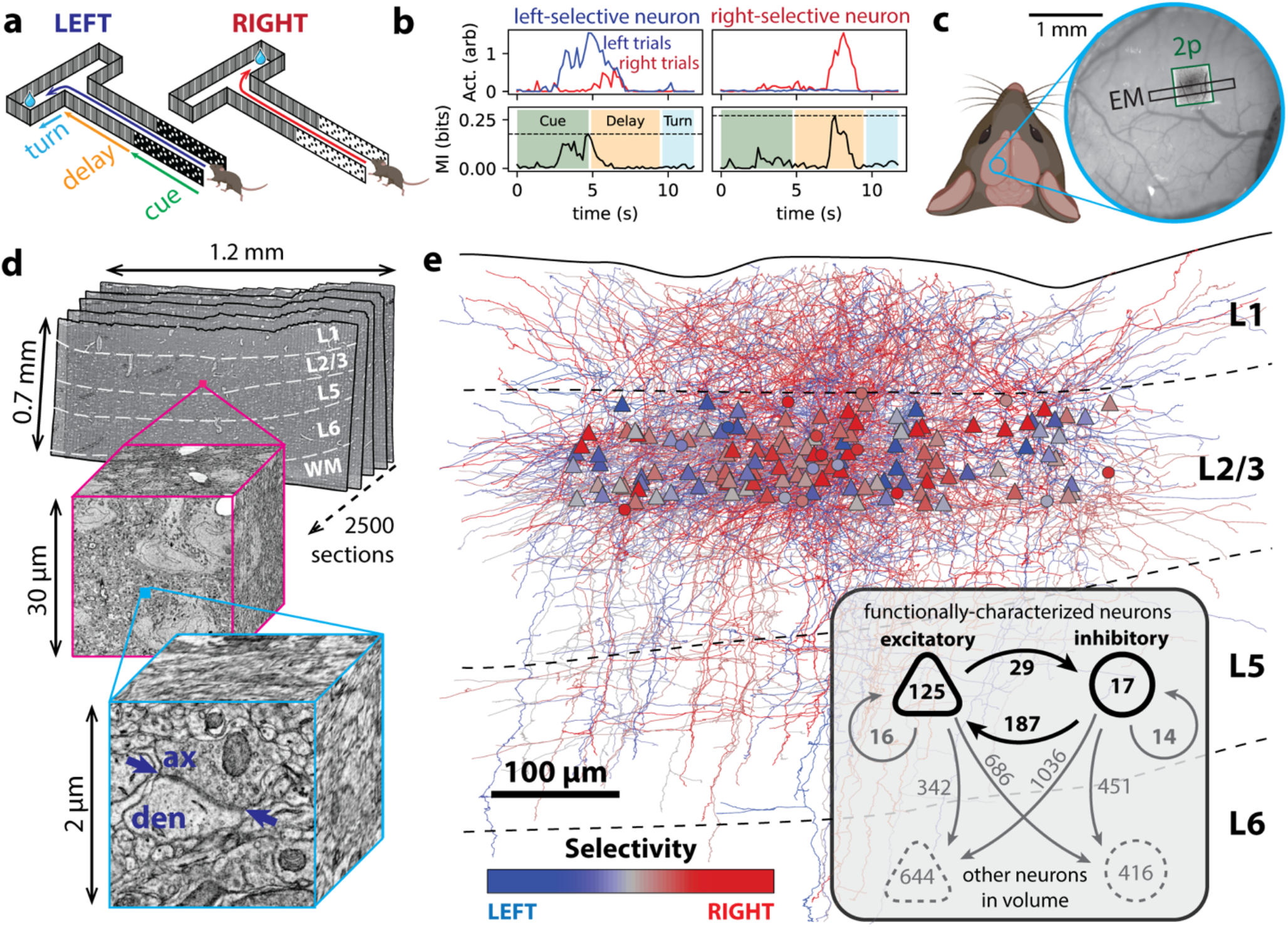
Behavior, functional imaging, and electron microscopy. **(a)** Schematic of decision-making behavior consisting of a navigational two-alternative forced-choice memory task performed in virtual reality. **(b)** Trial average activity for left (blue) and right (red) trials from two example selective neurons, plotted along with mutual information with trial type (bottom). The magnitude of the selectivity index was defined as the peak of the mutual information (dotted horizontal line). **(c)** Image of the cranial window, showing location of overlapping calcium imaging (green) and EM (black) datasets within PPC. **(d)** Schematic of volumetric EM dataset consisting of 2500 serial sections. Insets: images at progressively higher resolution, highlighting cell somata (magenta) and an individual synapse (cyan, arrows indicate PSD). **(e)** Reconstructed circuit in PPC, consisting of 125 excitatory (E, triangles) and 17 inhibitory (I, circles) neurons color coded by selectivity. Inset: summary of reconstructed circuit, indicating number of neurons of each type (within shapes) and number of synaptic connections (next to arrows). Direct E-to-I and I-to-E connections, which are analyzed in detail in Figs. 2 & 3, are shown in bold. ax – axon, den – dendrite, L – layer, WM – white matter.

We preserved the brain immediately after the conclusion of behavioral experiments and used EM to generate a high-resolution structural map of the same neurons whose activity was previously measured *in vivo* (Fig. 1c, Methods). We used the GridTape automated transmission EM pipeline^17^ to collect and image 2500 serial 40 nm thin sections and aligned them to form a 3D volume spanning all six cortical layers with ∼1.2 mm extent (medial-lateral) and ∼100 µm depth (anterior-posterior). This dataset encompasses approximately 0.1 mm^3^ at 4.3 × 4.3 × 40 nm per voxel resolution (Fig. 1d). We then co-registered the *in vivo* and EM data to match calcium imaging regions of interest to cell somata in the EM volume (Ext. Data Fig. 1e,f, Methods). Thus, we were able to relate behavior, neuronal activity, and high-resolution anatomy (including synaptic connectivity) for individual neurons in PPC.

Using the EM data, we reconstructed the axons and dendrites of the functionally characterized cells within the volume (Fig. 1e) and classified them as excitatory pyramidal cells or inhibitory interneurons based on their morphology (n = 125 pyramidal, 17 non-pyramidal, Supp. Fig. 2,3). Non-pyramidal cells in PPC were generally as selective as pyramidal cells (Ext. Data Fig. 1g, p = 0.37, K-S test), which is consistent with recent functional imaging experiments^10^. Selectivity of interneurons in PPC contrasts with V1, where interneurons are more broadly tuned to stimulus orientation than pyramidal cells^23,31,36–38^ (but see ^32–34^).

To map connectivity of the functionally characterized neurons, we annotated all of the outgoing synapses from their axons within the EM volume (2214 excitatory and 2418 inhibitory synapses) and traced the corresponding post-synaptic dendrites back to their somata (Fig, 1e inset). We also quantified the area of the post-synaptic densities (PSD areas) associated with each of the synapses (Fig. 1d inset, Methods), which correlates with functional synaptic strength^39^. To enable comparisons between PPC and V1, we also analyzed a previously generated EM volume of V1 containing layer 2/3 neurons with corresponding *in vivo* measurements of orientation tuning^26^ (Ext Data Fig. 1h,i).

**Ext. Data Fig. 1:**
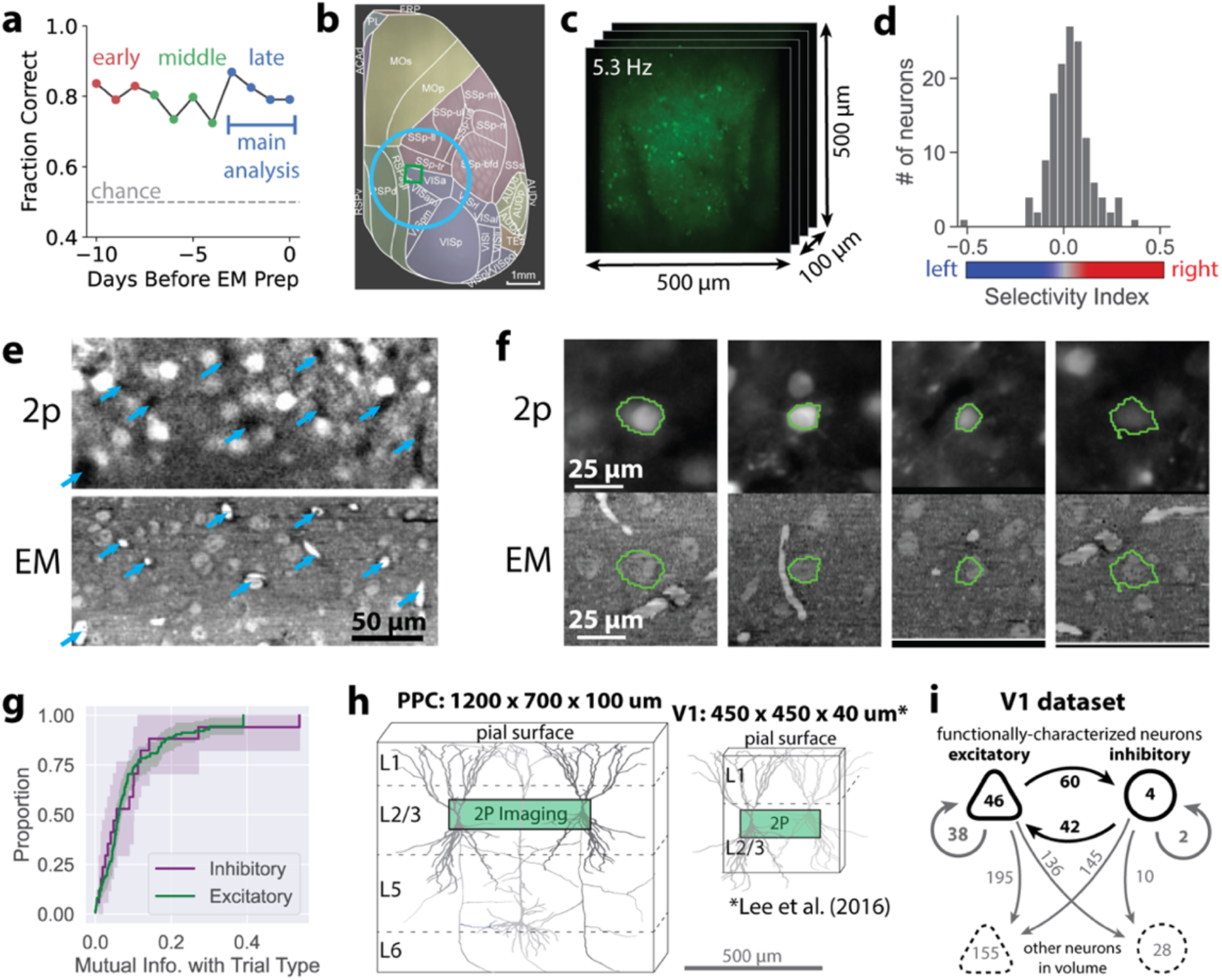
**(a)** Performance on the navigational two-alternative forced-choice memory task plotted over different daily sessions. Functional data from the last 4 sessions (“late”), which were closest in time to when the brain was preserved for EM, were used for most analyses. Earlier sessions (“early”, “middle”) were used for additional analyses investigating how structure-function relationships evolve over time (see Ext. Data Fig 2i-l, Ext. Data Fig. 3b-c). **(b)** Approximate location of cranial window and calcium imaging ROI relative to cortical regions (adapted from the Allen Mouse Brain Common Coordinate Framework^40^). Although this figure depicts the right hemisphere, the experimental data was collected from the left hemisphere. **(c)** Calcium imaging ROI: 4 500 µm x 500 µm planes in layer 2/3 (separated by 25 µm in z) were imaged at 5.3 Hz volume rate. **(d)** Selectivity index for functionally characterized neurons contained within the EM volume (n=142). **(e)** Corresponding slices from co-registered 2-photon calcium imaging (top) and EM (bottom) datasets. Cyan arrows – blood vessels used as landmarks. **(f)** Example cellular ROIs (green) co-registered to calcium imaging (top) and EM data (bottom). **(g)** Cumulative histograms of mutual information with trial type, demonstrating that selectivity was not statistically different between inhibitory (purple) and excitatory (green) neurons (p = 0.37, K-S test). Shading indicates 95% confidence intervals generated via bootstrap analysis. **(h)** EM volumes sizes for PPC and V1^26^. **(i)** Summary of reconstructed V1 connectivity, indicating number of neurons of each type (within shapes) and number of synaptic connections (next to arrows) among these neuron types.

### Selective E-to-I Connectivity

We first investigated how E-to-I connectivity in PPC is related to selective activity. Using reconstructed connectivity from EM, we estimated rates of excitatory connectivity with local inhibitory and excitatory partners (Fig. 2a, Methods). Local E-to-I connectivity rates were several times higher than E-to-E (Fig. 2b, E-to-E = 0.4±0.1%, E-to-I: = 1.7±0.1%, p<1e-14 Mann-Whitney U-test). This was largely due to preferential targeting of inhibitory neurons: 67% of excitatory synapses were E-to-I even though inhibitory neurons make up only about 20% of local neurons^41^ (Ext. Data Fig. 2a). Local excitatory connectivity rates were also lower in PPC than V1 (Ext. Data Fig. 2b), which was is surprising given that cortical association areas exhibit stronger functional coupling than sensory areas^42,43^. This difference likely arose because PPC pyramidal axons had a lower density of synapses (Ext. Data Fig. 2c-d), and their collaterals branched at right angles from their trunks, making them less likely to immediately ascend back to the local layer 2/3 circuit (Ext. Data Fig. 2e).

**Figure 2:**
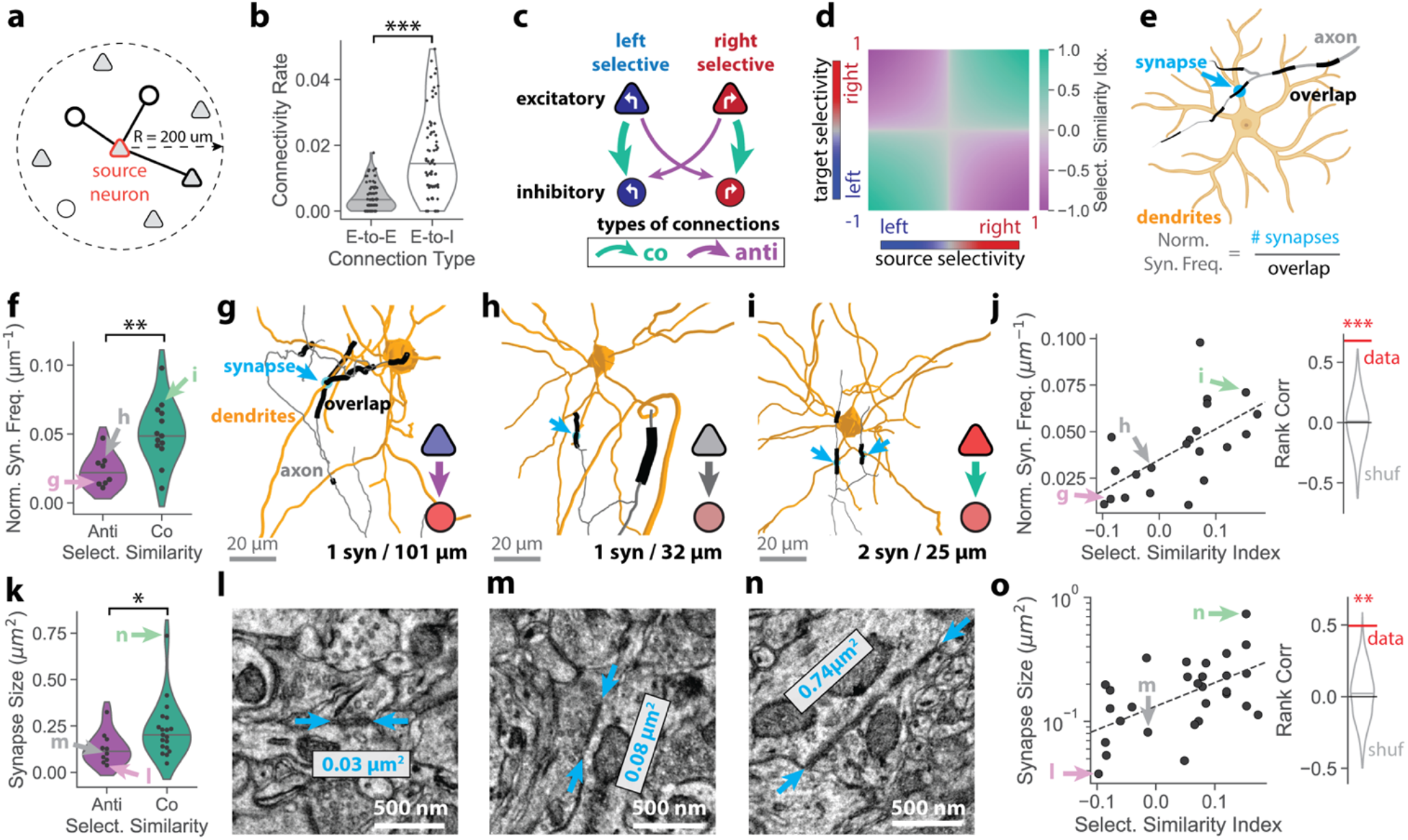
Co-selective Excitatory-to-Inhibitory Connectivity. **(a)** Local connectivity for a given source neuron (orange outline), is considered for neurons with cell bodies within 200 µm. The local connectivity rate is the proportion of excitatory (triangles) or inhibitory (circles) neurons with which the source neuron makes synaptic connections (lines). **(b)** Local E-to-I connectivity rates are several times higher than E-to-E in PPC (n = 70 source neurons, E-to-E: 0.4±0.1%, E-to-I: 1.7±0.1%, p < 1e-14, Mann-Whitney U test). **(c)** Schematic of E-to-I connections among functionally-characterized neurons. Connections are classified as co-selective (green) or anti-selective (purple). **(d)** Selectivity similarity index as a function of the selectivity index of the source and target neurons. Co-selective pairs have positive (green) and anti-selective pairs have negative values (purple). **(e)** Schematic of “normalized synapse frequency” metric which quantifies the likelihood of synapses between a specific axon-dendrite pair per µm of cable overlap (axon path length within 5 µm of the dendrite). **(f)** Normalized synapse frequency between co-selective (green) neurons are more than twice as frequent as anti-selective (purple) (anti = 0.024±0.004, n=8 connections; co: 0.051±0.006 µm^2^, n=13; p = 0.005, Mann-Whitney U test). **(g-i)** Example connections, including **(g)** strongly anti-selective, **(h)** weakly anti-selective, and **(i)** strongly co-selective pairs, colored as in (e). Gray – presynaptic axon, orange – post-synaptic dendrites, black – axon-dendrite overlap, cyan arrows – synaptic connections. Left/right selectivity of pre and post-synaptic neurons is indicated by colored icons. Number of synapses and total length of axon-dendrite overlaps (black) are indicated. Neuron pairs correspond to datapoints indicated by arrows in (f) and (j). **(j) Left:** Normalized synapse frequency is correlated to selectivity similarity index (n = 21 connections). Dotted line indicates linear fit. **Right:** Spearman’s rank correlation coefficient for data compared to random shuffles (c = 0.68, p < 0.001, permutation test). **(k)** Synapses between co-selective (green) neurons are almost twice as large as anti-selective (anti: 0.13±0.02, n=10 synapses; co: 0.23±0.03 µm^2^, n=19; p = 0.014, Mann-Whitney U test). **(l-n)** EM images showing example synapses between **(l)** strongly anti-selective, **(m)** weakly anti-selective, and **(n)** strongly co-selective neuron pairs. PSDs are indicated by cyan arrows and calculated PSD areas are shown by labels. Synapses correspond to datapoints indicated by arrows in (k) and (o) and are from the same connections shown in g,h,i, respectively. **(o**) **Left:** PSD area is correlated with selectivity similarity index (n = 29 synapses). Dotted line indicates linear fit. **Right:** Spearman’s rank correlation coefficient for data compared to random shuffles (c = 0.49, p = 0.002, permutation test). All statistics reported as mean ± standard error.

To determine if local E-to-I connectivity was functionally specific, we investigated if the selectivity of pre- and post-synaptic neurons predicted the likelihood of them forming synaptic connections. We hypothesized that the frequency of synaptic connections would be higher between neurons with the same selectivity (“co-selective”) than those with opposite selectivity (“anti-selective”) (Fig. 2c). To account for the distribution of selectivity strengths across neurons (Ext. Data. Fig. 1d), we defined a “selectivity similarity index”, which quantifies both how selective the pre- and post-synaptic neurons are and whether they are co- or anti-selective (Fig. 2d, Methods). Since the opportunity for neurons to make synaptic connections is limited to locations where their axons and dendrites come into close proximity, we normalized the number of synaptic connections by their axon/dendrite overlap^26,44^. This “normalized synapse frequency” measures the likelihood of connections between neurons independent of their locations and morphologies (Fig. 2e, Methods). We found that normalized synapse frequency was over two times higher for co-selective than anti-selective neuron pairs (Fig. 2f, anti = 0.024±0.004; co: 0.051±0.006 µm^2^; p = 0.005 Mann-Whitney U test). Furthermore, the highest and lowest synapse frequencies tended to be connections between strongly selective neurons (Fig. 2g-i). Indeed, the selectivity similarity index was strongly correlated with normalized synapse frequency (Fig. 2j, c = 0.68, p < 0.001, permutation test, Supp. Table 1). In contrast, in V1 we did not find a significant difference in normalized synapse density between co- and anti-selective E-to-I pairs (<45° or >45° difference in orientation tuning, respectively) (Ext. Data Fig. 2f,g), consistent with random E-to-I connectivity^23,31^.

We next asked if the strength of individual synapses was correlated with selectivity. Previously, it has been shown that the size of cortical excitatory synapses (PSD area) correlates with functional strength^39^. Here, we found that co-selective E-to-I synapses were nearly 2 times larger than anti-selective (Fig. 2k-n, anti: 0.13±0.02 µm^2^; co: 0.23±0.03 µm^2^; p = 0.014 Mann-Whitney U test). Synapse size was also strongly correlated with selectivity similarity index (Fig. 2o, c = 0.49, p = 0.002, permutation test). In contrast, E-to-I synapses between co-selective neurons in V1 were not larger than anti-selective neurons (Ext. Data Fig. 2h), providing additional evidence that selective connectivity differs between PPC and V1.

Given that the functional selectivity of PPC neurons can change over time periods of a few days^35^, we hypothesized that the observed structure-function relationships would be stronger for trials closer to when the brain was preserved. Comparing earlier with later behavioral sessions (Ext. Data Fig. 1a) revealed that structure-function correlations become weaker as one looks further back in time (Ext. Data Fig. 2i-l), suggesting that synaptic connections in PPC may also change over timescales of days.

**Extended Data Figure 2:**
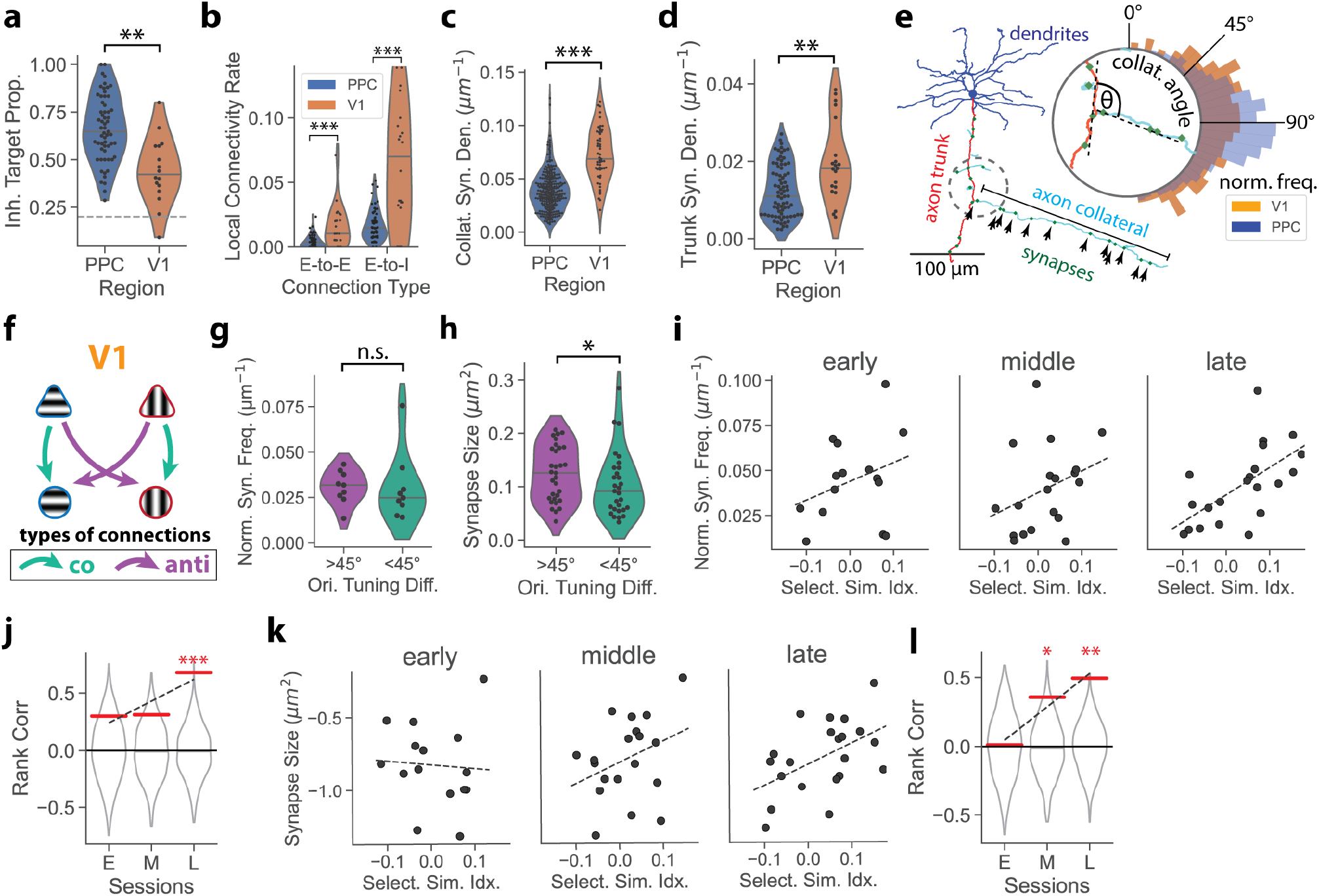
**(a)** Proportion of excitatory synapses that target inhibitory neurons is higher in PPC than V1 (PPC: 66 ± 2%, n = 59 neurons; V1: 44 ± 4%, n=16 neurons, p<1e-4, Mann-Whitney U test) and higher than the overall proportion of inhibitory neurons (∼20%, dashed line). **(b)** Local excitatory connectivity rates are lower in PPC than V1 for both E-to-E (PPC: 0.4 ± 0.1%, n = 73 source neurons; V1: 1.6 ± 0.3%, n = 22, p < 0.001, Mann-Whitney U test) and E-to-I connections (PPC: 1.7 ± 0.1%, n = 73; V1: 7.0 ± 1.2%, n = 22, p < 1e-4, Mann-Whitney U-test). **(c)** V1 axon collaterals have higher synapse density compared to PPC (PPC: 0.036 ± 0.001 syn/µm, n = 260 collaterals; V1 = 0.069 ± 0.004 syn/µm, n = 56, p < 1e-16, Mann-Whitney U test). **(d)** V1 axon trunks have higher synapse density compared to PPC (PPC: 0.012±0.001 syn/µm, n = 78 neurons; V1: 0.020±0.002 syn/µm, n = 21, p = 0.001, Mann-Whitney U test). **(e)** Example morphology of a pyramidal neuron, showing dendrites (blue), descending axon trunk (red), axon collaterals (green) and outgoing synapses (green). **Inset:** Initial angles for pyramidal axon collaterals are clustered near 90° in PPC, whereas V1 collaterals sample a broader range, including smaller angles that allow collaterals to ascend immediately back towards local layer 2/3 (n = 811, 112 collaterals for PPC, V1; p=.007, K-S test). **(f)** Schematic of E-to-I connections among functionally-characterized neurons in V1. Connections are classified as co-selective (<45° difference in peak orientation, green) or anti-selective (>45°, purple). **(g)** Normalized synapse frequencies are not significantly different for anti- (purple) and co-tuned (green) E-to-I connections in V1 (anti = 0.031±0.003, n=9 connections; co: 0.030±0.006 µm^2^, n=9; p = 0.18, Mann-Whitney U test). **(h)** Synapse size for co-selective (green) E-to-I connections are slightly *smaller* than anti-selective (purple) in V1 (anti: 0.12±0.009, n=31 synapses; co: 0.10±0.01 µm^2^, n=29; p = 0.03, Mann-Whitney U-test). **(i)** Correlations between normalized synapse frequency and similarity index for E-to-I connections in PPC, calculated using functional data from early (10-8 days before sacrifice), middle (7-4 days before), and late sessions (3-0 days before). **(j)** Spearman’s rank correlation coefficients corresponding to (i) are lower for earlier sessions (early: c = 0.30, n = 15 connections, p = 0.13, permutation test, middle: c = 0.31, n = 19, p = 0.09, late: c = 0.68, n = 21, p = .001), but the trend (positive slope) is not quite significant (p = 0.08, permutation test). Dotted line shows linear fit. **(k)** Correlations between PSD area and similarity index for E-to-I synapses in PPC, calculated for early, middle, and late sessions. **(l)** Spearman’s rank correlation coefficient corresponding to (k) are lower for earlier sessions (early: c = 0.01, n = 19 synapses, p = .49, permutation test, middle: c = 0.35, n = 24, p = .03, late: c = 0.49, n = 29, p = .003), and the trend (positive slope) is significant (p = 0.03, permutation test). Dotted line shows linear fit. All statistics reported as mean ± standard error.

### Selective I-to-E Connectivity

Having found functionally selective E-to-I connectivity in PPC, we next asked if I-to-E connectivity was also selective. Rates of local I-to-E connectivity were more than 3 times higher than E-to-I (Fig. 3a, I-to-E: 5.7±1.1%, E-to-I: 1.7±0.1, p < .001, Mann-Whitney U test), and inhibitory neurons were less biased towards targeting inhibitory post-synaptic partners compared to excitatory neurons (Ext. Data Fig. 3a).

**Fig. 3:**
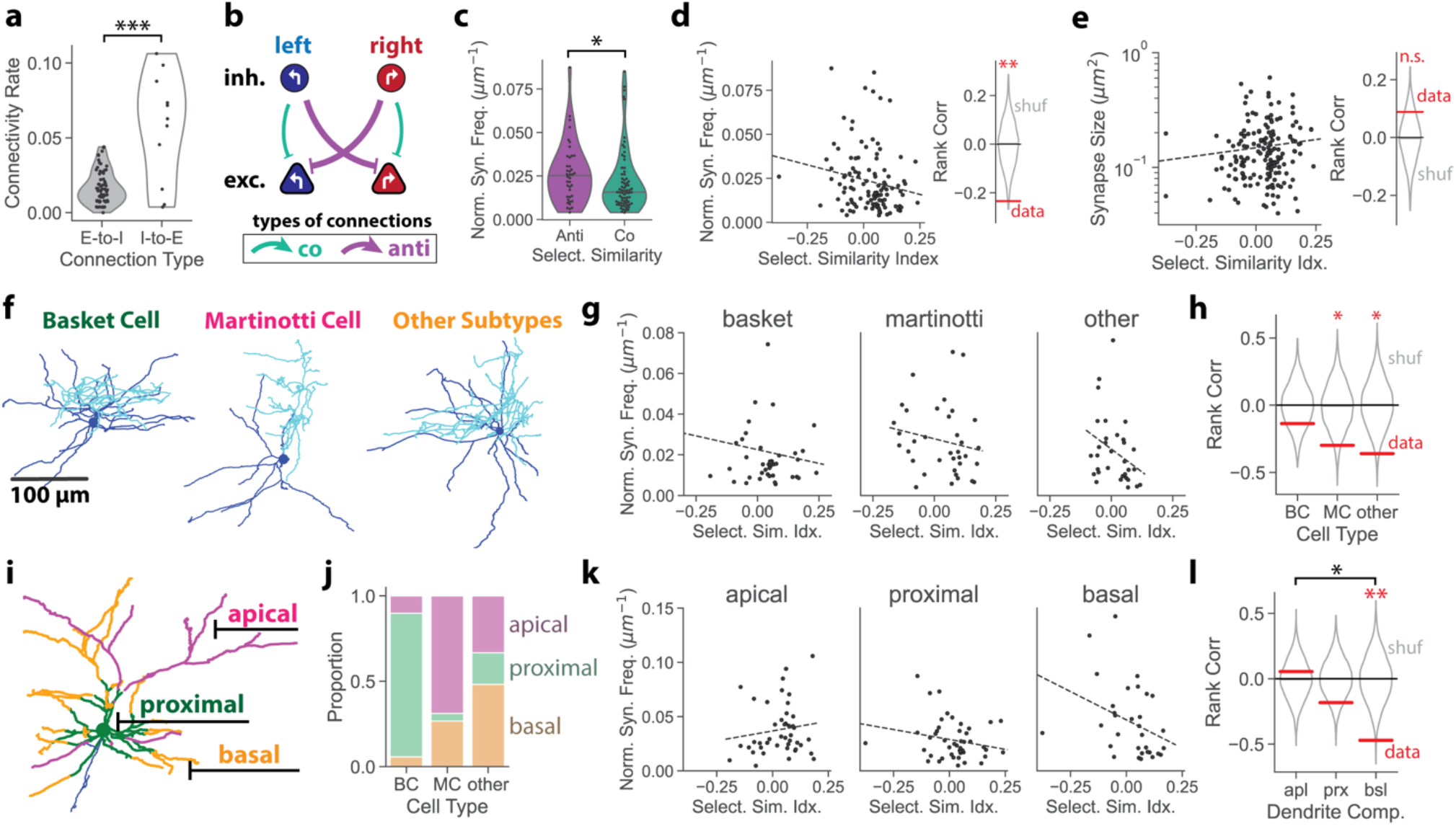
Anti-selective Inhibitory-to-Excitatory Connectivity. **(a)** Local I-to-E connectivity rates are higher than E-to-I (E-to-I: 1.7±0.1%, n = 70 source neurons, I-to-E: 5.7±1.1, n = 11, p < .001, Mann-Whitney U test). **(b)** Schematic of I-to-E connections among functionally-characterized neurons. Connections are classified as co- (green) or anti-selective (purple). **(c)** Normalized synapse frequency is greater for anti- (purple) than co-selective (green) I-to-E connections (anti: 0.027±0.003 syn/um, n=40 connections, co: 0.022±0.002 syn/um, n=71, p=0.01, Mann-Whitney U test). **(d) Left:** Normalized synapse frequency is negatively correlated with selectivity similarity index (n = 111 connections). Dotted line indicates linear fit. **Right:** Spearman’s rank correlation coefficient for data compared to random shuffles (c = −0.24, p = 0.004, permutation test). **(e) Left:** PSD area is not significantly correlated with selectivity similarity index (n = 169 synapses). Dotted line indicates linear fit. **Right:** Spearman’s rank correlation coefficient for data and random shuffles (c = 0.08, p = 0.13, permutation test). **(f)** Examples of basket cell (BC), Martinotti cell (MC), and other dendrite-targeting interneuron morphologies. Axons – cyan, dendrites – blue. **(g)** Normalized synapse frequency plotted as a function of selectivity similarity index for I-to-E connections from basket, Martinotti, and other interneuron subtypes. **(h)** Martinotti and other cell types have a significant negative correlation (basket: c=-0.14, n=43, p=0.19, permutation test, Martinotti: c=-0.30, n=36, p=0.04, other: c=-0.36, n=32, p = 0.02). Differences in correlation between cell types are not statistically significant. **(i)** Example pyramidal neuron with dendritic compartments labeled (apical – magenta, proximal – green, basal – orange, axon – blue). **(j)** Proportion of synapses made onto apical (pink, top), proximal (green, middle), and basal (orange, bottom) pyramidal dendrites for BC, MC, and other interneurons. Despite having a preferred target, all interneuron subtypes synapse onto a mix of dendritic compartments. **(k)** Normalized synapse frequency plotted as a function of selectivity similarity index for I-to-E connections targeting apical, proximal, and basal dendrites. **(l)** Basal-targeting connections have a significantly negative correlation (apical: c=0.05, n=44, p=0.33, proximal: c=-0.18, n=47, p=0.10, basal: c=-0.47, n=32, p =0.004), and basal dendrite connections are significantly more anti-selective than apical (p=0.04, permutation test with Bonferroni correction). All statistics reported as mean ± standard error.

To test if I-to-E connectivity is correlated with functional selectivity, we compared normalized synapse frequencies between co- and anti-selective neuron pairs (Fig. 3b). Normalized synapse frequency was slightly higher for anti-selective pairs (Fig. 3c, anti: 0.027±0.003 syn/um, co: 0.022±0.002 syn/um; p=0.01, Mann-Whitney U test), suggesting that inhibitory neurons preferentially target excitatory partners with opposite selectivity. Indeed, we found that the normalized synapse frequency of I-to-E connections was *negatively* correlated with selectivity similarity index (Fig. 3d: c = −0.23, p = 0.005, permutation test, Supp. Table 2). These correlations were weaker when using functional data from earlier sessions (Ext. Data Fig. 3b,c). In contrast to E-to-I connectivity, we did not find a significant correlation between the individual synapse sizes and selectivity similarity index (Fig. 3e, c = 0.08, p = 0.13, permutation test, cf. Fig. 2o), suggesting that selectivity I-to-E connectivity may be mediated more by number of synapses than the strength of individual synapses.

Cortical inhibitory interneurons consist of several genetically, physiologically, and morphologically distinct subtypes that can target specific compartments of pyramidal dendrites^45^. For example, basket cells (BCs) target cell somata and proximal dendrites, while Martinotti cells (MCs) project their axons upwards towards layer 1 where they target apical tufts^46^. We characterized inhibitory neurons as basket cells (BCs), Martinotti cells (MCs), or other dendrite-targeting cells based on their axon targeting (Fig. 3f, Ext. Data Fig. 3d,e, Methods) and compared their connectivity.

MCs and other dendrite-targeting cells exhibited a significant negative correlation between normalized synapse frequency and selectivity similarity, while BCs had weaker negative correlation (Fig. 3g,h: BC: c=-0.14, p=0.19, permutation test, MC: c=-0.30, p=0.04, other: c=-0.36, p=0.02). However, the differences in correlation coefficient between cell types were not significant.

Although the inhibitory subtypes each preferentially target a specific pyramidal dendrite compartment (Fig. 3i), they still make synapses onto multiple compartments (Fig. 3j). We hypothesized that connections targeting distinct dendrite compartments may exhibit different amounts of selectivity. To test this, we calculated normalized synapse frequencies for connections targeting apical, proximal, and basal dendrites separately, and quantified correlations with selectivity similarity index (Methods). We found that connections targeting basal dendrites were strongly anti-selective (Fig. 3k,l, c=-0.47, p =0.004, permutation test), and significantly more anti-selective than those targeting apical dendrites (p=0.04, permutation test with Bonferroni correction), which appeared largely unselective (c=0.04, p =0.33, permutation test). Connections targeting proximal dendrites were anti-selective, but not significantly so (Fig. 3k,l, c=-0.18, n=47, p=0.10), which is consistent with the overall weak anti-selectivity of basket cells (Fig. 3g,h). These results suggest that the anti-selectivity of I-to-E connections is driven primarily by synapses targeting basal dendrites and (to a lesser extent) proximal dendrites.

**Extended Data Figure 3:**
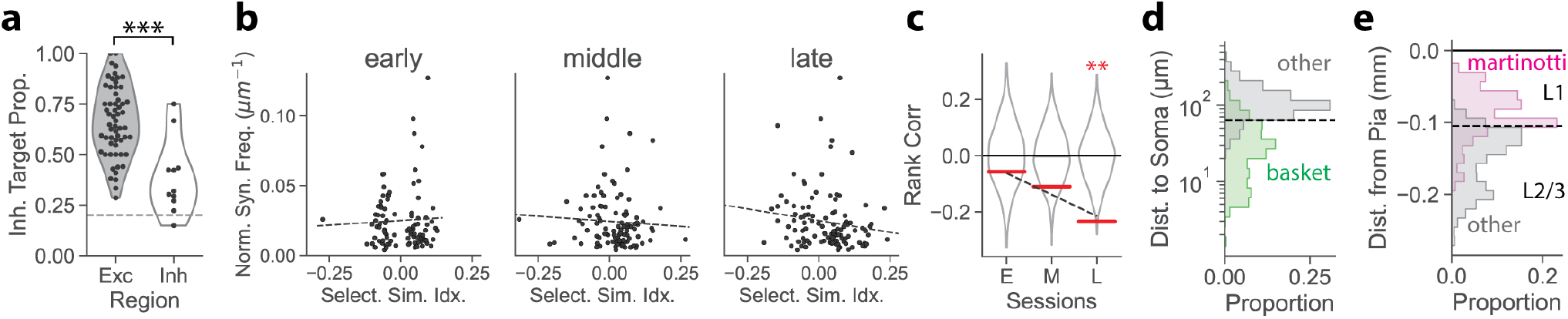
**(a)** PPC excitatory neurons target inhibitory partners at higher rates than inhibitory (excitatory: 66±2%, n=59 neurons, inhibitory: 38±5%, n=11, p < 10^−4^, Mann Whitney U test). Dotted line – estimated overall proportion of inhibitory neurons (∼20%)^41^. **(b)** Normalized synapse frequency plotted as a function of selectivity similarity index for I-to-E connections in PPC, calculated using functional data from early (10-8 days before sacrifice), middle (7-4 days before), and late (3-0 days before) sessions (early: n=93, middle: n=107, late: n=111). **(c)** Spearman’s rank correlation coefficients corresponding to (b) are less negative for earlier sessions (early: c=-0.06, p=0.29; middle: c=-0.11, p=0.12; late: c=-0.23, p=.004, permutation tests) but the trend (negative slope) is not quite significant (p = 0.09, permutation test). Dotted line shows linear fit. **(d)** Geodesic (along-the-dendrite) distances to post-synaptic soma from synapses made by basket cells (green, n=557 synapses) and other interneurons (gray, n=404). Basket cells preferentially target somata and proximal dendrites. Dotted line – threshold that maximally separates basket and non-basket synapses used to define proximal dendrites in further analysis (64 µm). **(e)** Depth (pia-to-white matter) of synapses from Martinotti (magenta, n = 255) and other interneurons (gray, n = 783). Martinotti axons ascend towards the pia and make synapses in layer 1. All statistics reported as mean ± standard error.

### Circuit Modeling

Together, selective E-to-I (Fig. 2) and I-to-E connectivity (Fig. 3) comprise an opponent inhibition motif (Fig. 4a, top). We used network modeling to investigate how opponent inhibition may support decision-making computations. We first studied a linear rate model^10,34,47,48^ comprising two excitatory and two inhibitory units. Left or right selective excitatory neurons (E_L_, E_R_) receive elevated external input during left or right trials and interact with left and right inhibitory neurons (I, I) (Fig. 4a, Methods). In networks with opponent inhibition, input onto E_L_ decreases E_R_ activity through feed-forward inhibition, which further amplifies E_L_ activity via feedback disinhibition^49–51^ (Fig 4b, Ext. Data Fig. 4b). In left trials, both suppression of E_R_ and amplification of E_L_ increased the distance between neural activity on left and right trials (Ext. Data Fig 4a, Methods), and this distance was thus larger for networks with stronger opponent inhibition (Ext. Data Fig 4b,c). As a consequence, networks with stronger opponent inhibition supported more accurate decoding of trial type in the presence of readout noise (Fig 4c,d, Ext. Data Fig 4d,e, Methods). This signal amplification through opponent inhibition occurs over a broad range of values of E-to-E selectivity, and even without recurrent excitatory connections (Supp. Fig. 4). When time-varying input noise was included, opponent inhibition amplified the signal more than it amplified the noise, therefore enhancing trial-type encoding (Supp. Fig. 5).

**Fig. 4:**
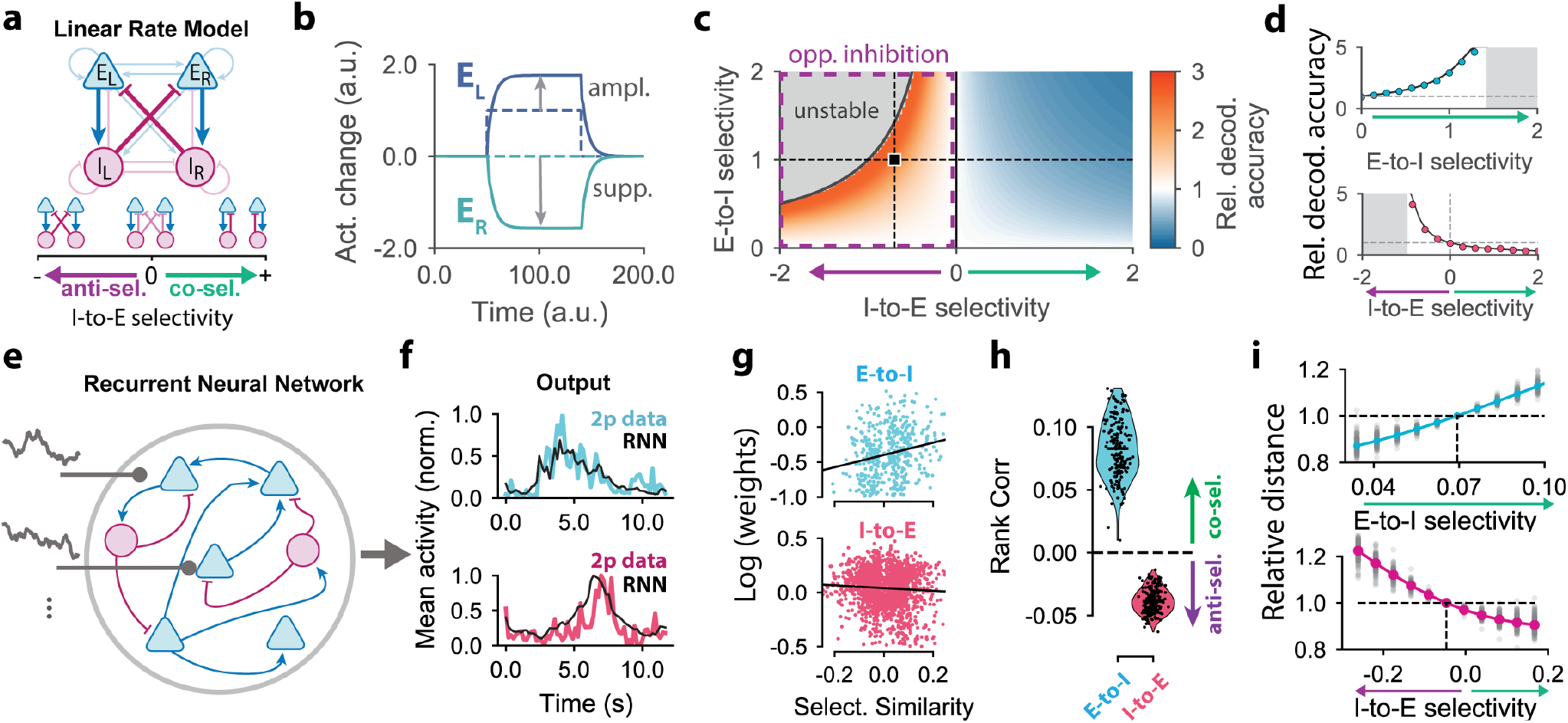
Opponent inhibition connectivity motif supports trial-type signal separation. **(a) Top:** Illustration of the linear rate model, comprising two excitatory and two inhibitory units. Left (or right) trial-type input is fed to the E_L_ (or E_R_) population. **Bottom:** Schematic of network variants in which I-to-E selectivity differs. Purple and green arrows indicate stronger anti- and co-selectivity, respectively. **(b)** Change in E_L_ and E_R_ activity in response to a left trial-type input for a network with opponent inhibition (co-selective E-to-I and anti-selective I-to-E). E_L_ is amplified and E_R_ is suppressed relative to the input (dotted lines). **(c)** Relative decoding accuracy, defined as the ratio of output to input decoding accuracy, as a function of E-to-I and I-to-E selectivity. The decoding accuracy is computed by linearly decoding the trial-type from the excitatory population in presence of readout noise. The black square denotes parameters used for panel (**b)**. The region with anti-selective I-to-E selectivity (purple arrow and box) corresponds to networks with opponent inhibition. The grey area corresponds to unstable network dynamics. **(d)** Relative decoding accuracy as a function of E-to-I (top) and I-to-E (bottom) selectivity, corresponding to two cuts of the phase plot (dashed lines in panel **(c)**). Purple and green arrows correspond to anti- and co-selectivity. **(e)** Illustration of a recurrent neural network (RNN) fit to the PPC population activity. The number of excitatory and inhibitory neurons in the RNN is matched to the experimental data. The networks are trained to reproduce trial-averaged PPC activity of matched neurons for left and right trials. **(f)** Examples of the PPC activity (colored lines) and RNN fits (black lines) for one excitatory (top) and one inhibitory (bottom) neuron. **(g)** Correlations between connectivity strength and selectivity similarity for E-to-I (top) and I-to-E (bottom) connections for a single RNN. E-to-I connections have a positive correlation (co-selective) while I-to-E have a negative correlation (anti-selective). **(h)** Spearman’s rank correlation coefficient between connection strengths and selectivity similarity for many trained RNNs (n=192). E-to-I connections are co-selective whereas I-to-E connections are anti-selective. **(i)** Normalized distance between left- and right RNN activity (averaged across time) as a function of selectivity perturbations. E-to-I (top) and I-to-E (bottom) connection weights were perturbed in a way that increases anti- (purple arrow) or co-selectivity (green arrows) without changing the average connection weight. The distance between trajectories is normalized by its value in the unperturbed network (dashed lines). Single networks and median values are shown by gray and colored dots, respectively (n=192).

While the linear rate model explains how opponent inhibition affects network coding, it does not include heterogeneity of connection weights nor capture the sequential dynamics observed in PPC^4,35^. To determine if its predictions hold for models incorporating these more realistic features, we built a recurrent neural network (RNN) model with the same number of excitatory and inhibitory neurons as the experimentally reconstructed circuit (Fig. 4e), and trained the connection weights of several individual RNNs to reproduce the measured calcium activity^52–54^. After training, the RNNs generated dynamics which accurately reproduced PPC activity^52^ (Fig. 4f). Although we did not place any constraints on the selectivity of the RNN connections, we found that the trained RNNs consistently exhibited co-selective E-to-I and anti-selective I-to-E motifs, similar to those found experimentally (Fig. 4g,h). To investigate if these motifs supported signal amplification, as predicted by the linear rate model, we systematically manipulated the RNN connectivity^55^ by perturbing the E-to-I or I-to-E selectivity around the trained values, and re-generated the dynamics using the new connections (Methods). Stronger opponent inhibition (stronger E-to-I co-selectivity or I-to-E anti-selectivity) amplified the separation between left and right population responses (Fig. 4i), further suggesting that opponent inhibition may enhance the coding of trial-type signals in PPC.

**Extended Data Figure 4:**
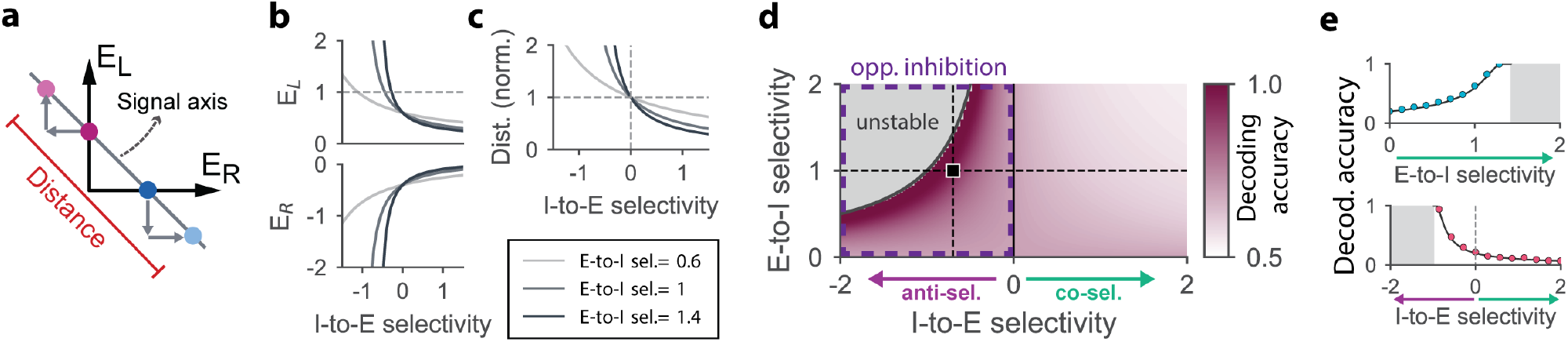
**(a)** Distance between mean responses to left and right trial types (colored dots) in the state-space of the excitatory neuron activity. This distance is enhanced by opponent inhibition between trial-type selective subnetworks. On left trial-types, both suppression of E_R_ and amplification of E_L_ (grey arrows) contribute to increased separation along the signal axis (and symmetrically for right trials). **(b)** Values of the steady state activity of E_L_ (top) and E_R_ (bottom) units as a function of the I-to-E (x-axis) and E-to-I (line colors) connection selectivity. **(c)** Normalized distance between mean activities corresponding to left and right trial-types (see panel **(a)**), as a function of I-to-E and E-to-I connection selectivity. **(d)** Decoding accuracy computed by linearly decoding the trial-type from the excitatory population in presence of readout noise, as a function of the E-to-I and I-to-E connection selectivity. The region with anti-selective I-to-E selectivity (purple arrow and box) corresponds to networks with opponent inhibition, the black square denotes the parameter values used in simulations of Fig 4b. The grey area corresponds to unstable network dynamics. **(e)** Decoding accuracy as a function of E-to-I (top) and I-to-E (bottom) selectivity, corresponding to two cuts of the phase plot (dashed lines in panel **(d)**). In all panels, purple and green arrows indicate the directions where connection motifs increase respectively their anti- and co-selectivity (see Fig. 4a).

## Discussion

### Functional Connectomic Dataset in PPC

We sought to understand relationships between trial-type selective neuron activity and synaptic connectivity in PPC. Although selective activity in PPC has been reported in many previous studies^4–11^, accompanying connectivity data has not been previously available. Here, we used automated serial-section transmission EM^17^ to image a volume in PPC with synapse resolution. Because neuronal arborizations extend over large distances in mammalian cortex, it is critical to image a large enough volume to sample them. The EM volume collected here in PPC contains a much larger volume than previous cortical EM datasets^23,25,26,30,56^ (but see ^29,57^), enabling reconstruction of substantial portions of axonal and dendritic arborizations, including synaptic connections made on distal branches. The resulting connectivity data, combined with behavioral and functional imaging data from the same animal, allowed us to reveal circuit motifs that support decision-making. Still, these connectivity motifs stem from a modest sample size of functionally-characterized neurons and synapses between them, which was constrained by the volume of the dataset. Future functional connectomic datasets involving much larger networks will likely reveal additional circuit motifs.

### Functionally Selective Connectivity Motifs

We found that the frequency and size of synaptic connections in PPC depended significantly on the selectivity of pre- and post-synaptic neurons. For E-to-I connections, co-selective synapses were larger and more frequent, whereas for I-to-E connections, anti-selective synapses were more frequent. We did not detect a difference in synapse size between co- and anti-selective I-to-E connections. However, synapse-size analysis for both E-to-I and I-to-E connections should be interpreted cautiously, as the correlation between synapse size and functional strength in cortex has only been directly measured for E-to-E synapses^39^.

The combination of co-selective E-to-I and anti-selective I-to-E comprises a competitive opponent inhibition motif, in which the activity of left-selective excitatory neurons suppresses the activity of right-selective ones, and vice versa. This motif has been shown to mediate action selection in zebrafish and *Drosophila*^58–60^, and a related motif has been proposed in ferret visual cortex^61^, but motifs of this type have not previously been found in cortical connectomes.

Previous work in mouse PPC proposed that selective connectivity motifs underlie choice-selective inhibitory activity, but could not rule out models with non-selective inhibition^10^. Here, the combination of neuronal activity measurements and EM-based connectomics in the same neurons has allowed identification of the underlying connectivity motifs.

Further analysis of I-to-E connectivity revealed that connections targeting basal dendrites were more selective than those targeting apical dendrites. This difference suggests that apical and basal dendrites can perform distinct computational roles^62^, and is consistent with the general idea that basal dendrites primarily receive local feedforward input while apical dendrites receive feedback signals^63,64^. However, we found that basket cells, Martinotti cells, and other dendrite-targeting interneurons were broadly similar in their overall selectivity, likely because they target a mix of dendrite types and the sample size of functionally-characterized interneurons was limited.

Selective inhibitory connectivity in PPC contrasts with V1, where previous connectomic analysis has suggested that E-to-I connectivity is non-selective in mice^23^ (but see ^65–67^). This suggests that specific inhibitory connectivity may be a distinct feature of PPC relative to V1, or alternatively is a consequence of plasticity induced by task training. These differences may underlie specialized functional roles of different cortical areas. Indeed, the selective inhibitory motifs we found in PPC promote separation of neural trajectories associated with competing behavioral choices, while primary sensory areas may privilege reliable encoding of diverse external stimuli. We also observed that E-to-E connectivity was sparser in PPC than in V1, which limited the sample size of E-to-E connections within the dataset and prevented us from confidently evaluating E-to-E selectivity in PPC. Sparse E-to-E connectivity in PPC is surprising given that V1 exhibits dense, like-to-like connectivity^26^, association areas exhibit stronger functional coupling between neurons^42,43^, and models of cortical decision-making often feature recurrent E-to-E connectivity^14^. It is possible that selective E-to-E connectivity may still occur over longer length-scales than the local circuit measured here. On the other hand, our circuit modelling suggests that decision-making computations can be achieved via opponent inhibition even with sparse or non-selective E-to-E connectivity (Supp. Fig. 4). Likewise, our dataset did not include enough functionally characterized inhibitory neurons to assess selectivity of I-to-I connectivity, but the effects of opponent inhibition are also robust over a wide range of values of I-to-I selectivity (Supp. Fig. 4).

### Decision-Making in Cortical Circuits

In models of decision-making, the formation of categorical choices is often facilitated by non-selective lateral inhibition^14^. Recently, it has been proposed that selective inhibition could play one of two possible roles: promoting competition with anti-selective I-to-E connectivity, or stabilizing dynamics through co-selective I-to-E connectivity^15^. These distinct contributions are also present in the linear rate model presented here (Supp. Fig. 4). Our anatomical data suggests that PPC lies in the competition regime and V1 lies in an intermediate regime characterized by non-selective I-to-E connectivity.

Although some decision-making models focus on the production of categorical choices via winner-take-all dynamics in nonlinear attractor models^15,47,48^, previous work suggests that during navigational tasks PPC produces more complex dynamics in which multiple activity patterns arise for each trial type^68^. These neural trajectories in PPC likely represent a wide range of task and behavioral variables, including the mouse’s choice, its navigational movements and position, and sensory cues from the environment^4,12,35,69–72^. For this reason, the model developed here focuses on a graded encoding of the choice signal, where the PPC circuit helps separate these multifaceted neural trajectories to enhance the encoding of the signals relevant for decision-making.

In summary, we discovered an anatomical opponent inhibition motif consisting of functionally selective connectivity between excitatory and inhibitory neurons in PPC. Using two complementary modeling approaches, we showed that this opponent inhibitory motif improves the encoding of trial-type information. Together, these results identify an anatomical connectivity motif in PPC that supports decision-making

## Acknowledgements

The authors thank Manuela Eroles, Karenna Ng, Lauren Hulshof, Triston Xie, Emily Dipietro, Trevor Khanna, Laurel Guo, Sarah Kushner, Elaina Phalen, Lia Decoursey, Elissa Zboinski, Leticia Sadilina, Mukul Narwani, Rholee Xu, Drasti Patel, Ziwei Wing Fan, Zach Diaks, Jimin Shin, Thedita Pedersen, and Amelia Buckner for neuron tracing; Xiuye Chen for providing image annotation software, Will Copeland, Joanna Kiperman, Benjamin Sanders, and Tobi Yusuf for assisting with EM imaging, Weiwei Lou, Jasper Phelps and Rui Zheng for assistance with image alignment; Amelia Buckner, Rholee Xu, and Seul Ah Kim for assistance with mouse training; Noah Petit for assistance with calcium imaging, Shin Kira, Alan Emanuel, and Alice Wang for providing mice; Greg Hood for providing alignment software (AlignTK); Philipp Schlegel for providing analysis software (pymaid and navis); Kenneth Hayworth for help with X-ray microCT processing; Mark Andermann, Rick Born, Tri Nguyen, Dianna Hidalgo and Daniel Wilson for comments on the manuscript. Some figures were created with BioRender.com

## Author Contributions

A.T.K., G.B., L.N.D., S.P, C.D.H., and W.-C.A.L. conceptualized the project and designed experiments. L.N.D. performed mouse behavior and calcium imaging experiments. W.-C.A.L., D.G.C.H., and A.T.K. prepared tissue samples for EM. D.G.C.H. and A.T.K. performed GridTape sectioning. B.J.G. and A.T.K. developed automated EM techniques and performed EM imaging. A.T.K. and L.A.T. performed EM image processing and alignment. M.K. and A.T.K. performed co-registration between calcium imaging and EM datasets. J.H. and A.T.K. performed and managed neuron tracing and data annotation. A.T.K performed analysis of circuit connectivity. G.B. performed circuit modeling and analysis. A.T.K., G.B., S.P., C.D.H., and W.-C.A.L. wrote the paper, and all authors assisted in reviewing and revising the manuscript.

## Data and Materials Availability

Data, software and analysis code will be made publicly available upon publication of the manuscript. In the interim, to request access please contact

