## Supplementary Material for "Synaptic wiring motifs in posterior parietal cortex support decision-making"

This document includes:

1. Methods (pg. 2)
2. Supplementary Tables (pg. 19)
3. Supplementary Figures (pg. 22)

### Methods

#### Behavior and calcium imaging

We trained mice to perform a two-alternative forced-choice delay task in a virtual reality T-maze and performed *in-vivo* calcium imaging as previously described[4,35] (Fig. 1a). Briefly, the task consisted of cue, delay, and turn phases. During the cue phase, mice were presented with one of two visual cues on the walls of the T-maze. In the second half of the T-maze (the delay period), the cues were replaced by a neural gray. At the T-intersection, mice turned left or right based on the presented cue to earn a water reward. Mice performed the task by running on a spherical treadmill and were presented with visual stimuli projected onto a screen using the ViRMEn software engine[73] (https://pni.princeton.edu/pni-software-tools/virmen).

We used raw calcium imaging and behavioral data from a mouse originally from a previous study[35]. Briefly, it was a male C57BL/6J mouse (The Jackson Laboratory) that was 8 weeks old at the start of behavioral training, 14-18 weeks old during imaging, and 18-19 weeks old when the brain was preserved for EM imaging. GCaMP6m was expressed in left-hemisphere PPC layer 2/3 neurons via viral injection (AAV2/1-synapsin-1-GCaMP6m) and neuronal activity was recorded during behavior using a 2-photon microscope (5.3 Hz volume imaging rate). Behavior and functional imaging were recorded every day for ~30 consecutive days (with a few 1-day breaks). However, for the main analysis in this work, we focused on data from the last 4 sessions. For additional analyses investigating how structure-function relationships evolve over time, we used sessions 10-8 days before sacrifice (early), 7-4 days before (middle) and the last 4 days (late) (see Ext. Data Fig. 1a, Ext. Data Fig. 2i-l, Ext. Data Fig. 3b-c).

For each session, we performed source extraction using Suite2p[74] (https://www.suite2p.org) and manually screened the resulting ROIs to obtain putative cell somata and associated calcium signals. The dF/F calcium signals were deconvolved using the constrained FOOPSI algorithm[75] (https://github.com/epnev/constrained-foopsi) to obtain an event rate that estimates the relative firing rate of each neuron over time. We synchronized the event rates to the behavioral trials structure to obtain trial-wise event rates for each neuron. Because the length of trials can vary based on the how quickly the mouse runs, we synchronized based on 3 landmarks during the trial: the beginning of the trial, the start of the delay period, and the end of the trial. Trial aligned event rates had 76 time points (each 188 ms) according to the following scheme:

| Time Points | Trial Epoch | Landmark Time Point |
|---|---|---|
| 1-13 | Inter-trial interval before | 14 – start of trial (running onset) |
| 14-26 | Cue (beginning) | 14 – start of trial (running onset) |
| 27-38 | Cue (end) | 39 – start of delay period (cue offset) |
| 39-51 | Delay | 39 – start of delay period (cue offset) |
| 52-64 | Turn | 64 – end of trial (reward given or omitted) |
| 65-76 | Inter-trial interval after | 64 – end of trial (reward given or omitted) |

Thus, synchronized trials were not completely continuous in time (there are 2 discontinuities). We then calculated trial-average event rates by averaging over left and right trials separately. For selectivity analysis, only time points after the beginning of the trial (time points 14-76) were used.

#### Information-theoretic selectivity index

For each (putative) neuron in each session, we calculated an information-theoretic choice-selectivity index in the following way. The event rates were converted to binary by setting all non-zero values to 1. This binarized event rate can be interpreted as a measure of when the neuron is active versus inactive. Then, as done previously[42], we calculated the instantaneous mutual information (MI)[76] between this binarized event rate and the identity of the sensory cue (which has 2 possible values for right and left trials) for each time-point in the trial using the Neuroscience Information Theory Toolbox[77]. Thus, the definition of "trial type" can be understood as the type of cue presented to the mouse.  All left and right-turn trials were included in analysis, including error trials in which the mouse turns the wrong direction. Alternative analyses, such as including only correct trials or defining the trial type based on the mouse's choice, lead to slightly different values of selectivity index, but did not significantly change the results of this study. We did not apply a finite sampling bias correction[78] because the bias was expected to be negligible given the trial numerosity and its application would have increased the variance of MI estimation. The MI has units of bits and is bounded between 0 and 1. We defined the magnitude of the selectivity index to be the maximum value of the MI across time points. The time point of peak MI was different for different neurons. We defined the sign of the selectivity index based on the trial-average event rates at the timepoint of peak MI: the selectivity index was given a negative sign if the trial-average binarized event rate was greater on left trials than right. Thus, the selectivity index ranges from -1 to +1, with the sign indicating preferred trial type and the magnitude indicating MI with trial type. Selectivity indices were calculated separately for each behavioral session. To obtain the overall selectivity index for each neuron, the selectivity index was averaged across the last 4 behavioral sessions before the mouse was sacrificed (see "Correspondence between in-vivo and EM data" below for alignment of multiple sessions and EM data). For additional analyses investigating how structure-function relationships evolve over time, we calculated selectivity index separately for "early", "middle" and "late" sessions (see Ext. Data Fig. 1a, Ext. Data Fig. 2i-l, Ext. Data Fig. 3b-c).

This information-theoretic definition of selectivity index is distinct from metrics based on receiver-operator characteristic analysis[10] or trial-averaged activity[4,35] used in other studies. In contrast to these other metrics, which are based on trial-average activity, this MI metric quantifies information available in individual trials, is robust to trial-to-trial variability, can capture non-linear tuning, and is more specific to the time-points during the trial where selectivity is highest. However, these different definitions of selectivity index are highly correlated to each other, the main conclusions of this study also hold when using alternative definitions of selectivity index.

It is worth noting in our dataset the number of right-selective neurons outnumbered the left-selective (Ext. Data Fig. 1d, Supp. Fig. 1). As a result, the number of functionally-characterized left-selective inhibitory neurons was small. Thus, our findings on choice-selective connectivity primarily comes from connections involving left-selective inhibitory neurons.

#### EM Dataset

On the last day of *in-vivo* imaging, we injected the tail vein with a fluorescent dye to label blood vessels (rhodamine B isothiocyanate–Dextran (MW ~70k), 5% v/v, Sigma) and acquired an anatomical 2p reference stack of the imaging ROI in PPC (green channel - GCaMP6m for cells, red channel – rhodamine for blood vessels; Ext. Data. Fig. 1e). After performing behavior and calcium imaging, the animal was perfused transcardially (2% formaldehyde/2.5% glutaraldehyde in 0.1 M cacodylate buffer with 0.04% $CaCl2$ ) and the brain was prepared for EM imaging as previously described[26] . Briefly, 200 μm thick coronal vibratome sections were cut, post-fixed, and en bloc stained with 1% osmium tetroxide/1.5% potassium ferrocyanide followed by 1% uranyl acetate, dehydrated with a graded ethanol series, and embedded in resin (TAAB 812 Epon, Canemco).

We cut serial 1 μm thick sections from regions a few sections away from PPC, stained them with toluidine blue (EMS), imaged them with light microscopy, and aligned them to large blood vessels on the surface of the brain (from photos taken through the cranial window) to estimate which sections overlap with the PPC regions imaged *in vivo*. Then, we used micro-CT (Zeiss Versa) to confirm the correct vibratome section by registering corresponding vasculature between the micro-CT volume and 2p reference stack.

We cut a series of 2500 thin-sections (40-45 nm thick) and imaged them using the GridTape system as previously described[17]. Briefly, we trimmed the tissue block to mesa containing the tissue of interest using an ultramicrotome (Leica UC7) and a diamond trimming knife (EMS-Diatome). Then, using an automated tape ultramicrotome (ATUM), we sectioned and collected 2500 sections onto a reel of GridTape over a period of about 15 hours. During pickup, 37 sections (out of 2500, comprising 1.4%) were did not adhere to the transparent slots on the GridTape and 8 sections (0.3%) had ruptured film supports.

Following sectioning and pickup, we post-stained the sections with lead citrate using a semi-automated reel-to-reel system. The sections were imaged using automated transmission EM system over a period of about 6 months. For each section, an ROI approximately 1.2 x 0.7 mm was imaged (for the first 800 sections, a larger ROI of about 1.5 x 0.7 mm was imaged). During staining and imaging, 38 (1.5%) additional sections were damaged in a way that precluded successful imaging. Overall, 2427 (97.1%) of the intended 2500 sections were successfully imaged. The missed sections included 1 4-section gap, 2 3-section gaps, and 10 2-section gaps. The dataset consisted of ~400 TB of raw data (16-bit images). The raw images were converted to 8-bit format and stitched together to form a contiguous 3D volume using an elastic spring mesh algorithm (AlignTK).

#### Co-registration between *in-vivo* and EM data

On the last day of *in-vivo* imaging, we recorded a volumetric reference stack of the ROI in PPC with $(1\ \mu m)^3$ voxel size, which was used as a bridge between the calcium imaging planes for each session (4 planes separated by 25 µm in z) and the EM volume. First, we co-registered the reference stack to a down-sampled version of the EM volume (Ext. Data Fig. 1e). Then, for each session, we co-registered the imaging planes to the reference stack. Both registrations were calculated by manually identifying a moderate number of correspondence points (~10-30) using the ImageJ plugin BigWarp[79] (https://imagej.net/BigWarp) to calculate an affine transformation matrix (custom MATLAB code).

Using these affine transformations, we overlaid the extracted source ROIs from each calcium imaging session onto the EM space, and manually inspected each ROI for matching cell somata in the EM volume (Ext. Data Fig. 1f). Some ROIs were not associated with any somata, presumably because they were other objects such as large dendrites. To increase confidence, we matched ROIs to EM for multiple sessions simultaneously, and used the trial-aligned activity to help determine if ROIs in different sessions were from the same neuron. Activity was usually – but not always – similar for the same neuron over several sessions. At the conclusion of this correspondence process, we identified 142 functionally characterized neurons within the EM volume with matching calcium imaging ROIs. For most neurons, matching ROIs were found in multiple – but not all – sessions. This might be because they were not identified by the source extraction algorithm, or because they were excluded due to uncertainty in the co-registration and manual matching procedure.

#### Neuronal Circuit Reconstruction

The morphology and connectivity of the functionally characterized neurons were reconstructed via manual tracing by a team of annotators using the CATMAID[23,26,80,81] collaborative annotation software. Starting from the soma (which was previously co-registered to calcium imaging source ROIs), all branches (including axons and dendrites) were traced completely until they either ended or reached the boundary of the EM volume. In some cases, data quality issues such as missing sections or poor image quality prevented further tracing. The functionally characterized neurons were classified as pyramidal or non-pyramidal (inhibitory) based on their reconstructed morphology. Pyramidal cells were classified as such based on characteristic features including a prominent, pial-projecting apical dendrite, outward/downward projecting basal dendrites, and downward-projecting axon (Supp. Fig. 1). Non-pyramidal cells had a variety of morphologies likely corresponding to distinct interneuron subtypes (Supp. Fig. 2)

Synaptic connections were identified by characteristic ultrastructural features[82] including concentrated synaptic vesicles and a darkening/thickening of the membrane between the pre- and post-synaptic compartments (post-synaptic density). We annotated synapses in a way that also encodes an estimate of the area of the post-synaptic density (PSD). Connector objects were annotated on the section in which the PSD appeared longest, and the pre- and post-synaptic nodes were placed in a way such that the distance between them was equal to the length of the PSD (Fig. 1d, Fig. 2l,m,n). This length was taken to be the diameter of circular PSD to estimate PSD area. Although PSDs are not exactly circular, the errors inherent to this estimation are likely small compared to the large variations in size from synapse to synapse[28,83].

Starting from the outgoing synapses on the axons of functionally characterized neurons, each post-synaptic neuron was traced until either the soma was found or the neuron left the edge of the EM volume. Those post-synaptic partners with somas within the volume were classified as pyramidal or non-pyramidal as described above. Some of the post-synaptic partners were other functionally characterized neurons (direct connections). Thus, the first-order output connectivity of the functionally characterized neurons was traced completely within the EM volume (Fig. 1e, inset).

All tracing was reviewed a second time by an independent reviewer. In general, we took a conservative approach to terminate ambiguous continuations to avoid merge errors.

#### Primary Visual Cortex Dataset (V1)

We used a primary visual cortex (V1) EM dataset, neuronal circuit reconstruction, and associated neuron activity data that was generated as part of a previous study[26]. Briefly, this dataset included neuronal morphology, connectivity, and functional characteristics (orientation tuning and preferred orientation, derived from calcium imaging) (Ext. Data Fig. 1h). We performed some additional neuron tracing and annotation beyond what was previously completed. To quantify similarity of functional activity, we defined an orientation-similarity metric analogous to choice-similarity for PPC.

$$s = \sqrt{c_1 c_2}(45° - \Delta\theta)$$

Where $c_1$ and $c_2$ are the orientation tuning for each neuron (1- circular variance) defined on $c \in [0,1]$, and $\Delta\theta$ is the difference in peak orientation defined on $\Delta\theta \in [0°, 90°]$. Thus, $s \in [0,1]$ and encodes both how sharply and how similarly tuned the two neurons are.

#### Connectivity Analysis

Analysis of neuron morphology and connectivity was performed by querying the CATMAID database and performing calculations using custom python code based on the navis (https://github.com/navis-org/navis) and pymaid (https://github.com/navis-org/pymaid) libraries. Specific details for particular analyses are provide below.

**Local connectivity rates** (Fig. 2a,b; Ext. Data Fig. 2b, Fig. 3a)

For each functionally characterized neuron (source neuron), all annotated neurons with somas within 200 µm of the source neuron soma were considered potential local partners. Potential partners were not limited to other functionally-characterized neurons. Since we were interested in cell-type-specific connectivity, neurons for which we were unable to classify a cell type (pyramidal or non-pyramidal) were excluded. The connectivity rates reported are the number of connected partners divided by the number of potential partners.

**Inhibitory target proportion** (Ext. Data. Fig. 2a, Ext. Data Fig. 3a)

For each functionally characterized neuron (source neuron), the inhibitory target proportion was the number of synapses that target inhibitory post-synaptic partners (PSPs) divided by the total number of PSPs. Since we were interested in cell-type-specific connectivity, PSPs for which we were unable to classify a cell type (pyramidal or non-pyramidal) were excluded.

**Axon synapse densities** (Ext. Data Fig. 2c,d)

Each pyramidal axon was divided into the trunk (the portion that extends from the soma downwards towards the white matter) and collaterals (branches that come off of the trunk). The collateral synapse density (Ext. Data Fig. 2c) was the number of synapses on the collateral divided by the total path length of the collateral. The trunk synapse (Ext. Data Fig. 2d) density was the number of synapses on the trunk divided by the path length of the trunk.

**Pyramidal axon collateral angle** (Ext. Data Fig. 2e)

For each axon collateral, the collateral angle vector was defined to point from where the collateral branches off the trunk to where it ends up after 20 um of pathlength. The collateral angle was the angle between this vector and the vertical direction (WM-pia axis). Axon trunks were generally close to vertical (within ~10º).

**Structure-function analyses** (Fig. 2c-o, Ext. Data Fig. 2f-l, Fig 3d-e,g-h,k-l, Ext. Data Fig. 3b-c)

These analyses quantified relationships between functional selectivity and connectivity for pairs of connected neurons. A pair of neurons was considered "co-selective" if their selectivity indices (see "Information-theoretic selectivity index" above) had the same size, and "anti-selective" for opposite sign (Fig. 2c,k, Ext. Data 2f,g,h, Fig. 3b,c). To quantify in a continuous manner how similar the selectivity is between two neurons, we defined a "choice similarity index" $s$ (Fig. 2d):

$$s = \pm\sqrt{|c_1||c_2|}$$

where $c_1$ and $c_2$ are the selectivity indices for the two neurons, and the sign of $s$ is positive if $c_1$ and $c_2$ have the same sign (co-selective) and negative if they have opposite sign (anti-selective). Thus, the pair similarity index encodes both directionality and strength of selectivity.

In these analyses we calculated correlations between pair similarity index and anatomical connectivity features: normalized synapse frequency or PSD area. We defined the normalized synapse frequency $f$ between a source and target neuron:

$$f = N/L_a$$

where $N$ is the number of synaptic connections and $L_a$ is the potential synapse length. $L_a$ is the path length of the source neuron's axon that is within 5 um (maximal spine length) of dendrites of the target neuron (Fig. 2e). $L_a$ quantifies how many opportunities there are for the two neurons to connect based on their fine-scale morphology[26]. The PSD area was calculated for each individual synaptic connection based on synapse annotations which estimated the PSD size (see "Neuronal Circuit Reconstruction" above). The PSD area has been shown to be a correlate of synapse strength (at least for E-to-E synapses in sensory cortex[39]).

We calculated Spearman's rank correlation between the functional (pair selectivity) and anatomical (pair synapse density or PSD area) measures, and performed permutation tests to assess the significance of any correlations. For pair synapse density, each datapoint was a connection between two neurons, which can involve one or more synapses. For PSD area, each datapoint was a single synapse. Spearman's rank correlation was chosen over Pearson linear correlation because it is less susceptible to outliers and more applicable for non-linear relationships. To generate a null distribution, we shuffled the selectivity indices among the source and target neurons, then recalculated the choice similarity based on these shuffled selectivity indices. This effectively removes structure-function relationships in the data. The p-value calculated with the permutation test was the proportion of correlation coefficients from the shuffle distribution that were greater than the measured correlation coefficient (or less than, if the coefficient was negative).

**Early, middle and late behavioral sessions** (Ext. Data Fig. 1a, Ext. Data Fig. 2i-l, Ext. Data Fig. 3b-c)

For analyses investigating how structure-function relationships evolve over time, functional selectivity indices were calculated separately from early (10-8 days before sacrifice), middle (7-4 days before), and late (3-0 days before) sessions. Correlations between similarity indices and connectivity (normalized synapse frequency or PSD area) were then calculated separately for early, middle, and late sessions.

The significance of the trend in correlation coefficients across early, middle, and late sessions was calculated using a permutation test. The trend was quantified by the slope of a linear fit to the correlation coefficients across early, middle, and late sessions. To create the null distribution, the early/middle/late labels for all data points were shuffled, correlation coefficients for each session group were recalculated, and the slopes of the linear fits were recorded. The p-value calculated with the permutation test was the proportion of the shuffled slopes that were greater than the slope of the data (or less than, if the slope was negative).

**Interneuron subtypes: basket, Martinotti, and other cell types** (Fig. 3f,g,h,j, Ext. Data Fig. 3d,e)

Functionally-characterized inhibitory neurons were classified by their axon targeting. Neurons that made a significant number of synapses onto post-synaptic somata (<20 μm from the soma) were classified as basket cells (Fig. 3f,g,h, Ext. Data. Fig. 3d). Neurons whose axons projected towards the pial surface and made a significant number of synapses in layer 1 were classified as Martinotti cells (Fig. 3f,g,h, Ext Data. Fig. 3e). Neurons that were not classified as basket or Martinotti cells were considered "other" cell types. Although it is generally established that cortical basket cells express parvalbumin (PV+) and Martinotti cells express somatostatin (SST+), our experiments did not include confirmation of the genetic identity of these cells.

**Proximal, apical, and basal dendrites** (Fig. 3i-l)

Dendrites of functionally-characterized excitatory neurons were classified as proximal, apical, or basal. Dendrites within a distance d = 64 μm (pathlength or "along-the-arbor distance) were considered proximal. This distance threshold calculated from the distribution of I-to-E synapses – it was the value that maximally separated basket cell synapses from non-basket cell synapses. Specifically, the chosen value of d maximized the sum of the proportion of basket cell synapses made onto proximal dendrites and the proportion of non-basket cell synapses made onto non-proximal dendrites (see Ext. Data Fig. 3d).

Apical dendrites were identified by a pial-projecting apical trunk branches extensively in layer 1. All branches downstream of the apical trunk were considered apical, even branches that emerged early and ramified in layer 2/3. All other (non-apical) dendrites were classified as basal dendrites. The portion of basal and apical dendrites within 64 μm of the soma were all considered proximal dendrites.

To calculate normalized synapse frequency for proximal, apical, and basal dendrites, only synapses made onto the specific dendrites were included in the numerator (N), and only cable overlap with the specific dendrites were included in the denominator ($L_a$).

**Comparisons of structure-function correlation coefficients** (Fig. 3g-h,k-l)

To assess the significance of differences between correlation coefficients calculated for multiple conditions (e.g. basket/Martinotti/other cell types, or proximal/apical/basal dendrites) we performed a permutation test. To create the null distribution, the condition types for the data points were shuffled, the correlation coefficients for each condition were recalculated, and the differences between correlation coefficients across conditions were recalculated. The p-value calculated for each difference between conditions was the proportion of differences from the shuffle distribution that were greater than the differences in the data (or less than, if the difference was negative). The p-values were adjusted by multiplying by 3 (the number of possible comparisons, in these cases $m = {}_3C_2 = 3$) to account for multiple comparisons testing (Bonferonni correction).

#### Network models

##### The linear rate model

To study how different connectivity motifs may impact PPC function, we examine a simple linear network model with recurrent connections. The network comprises excitatory and inhibitory units organized in a left and a right trial-type-selective subnetworks, each of them including one excitatory and one inhibitory unit (respectively $E_L, I_L$ and $E_R, I_R$ for left and right subnetworks). The dynamics of the network is defined by the differential equation:

$$\dot{r}_i = -r_i + \sum_{j=1}^{N=4} J_{ij} r_j + I_{ext,i} + \eta_i(t), \tag{1}$$

where $\mathbf{r} = \left(r_{E_L}, r_{I_L}, r_{E_R}, r_{I_R}\right)$ represents the firing rate deviation from the baseline activity level (here corresponding to $r_i = 0$ for all neurons $i$). In the linear regime, the baseline activity can be modulated by constant external inputs to the units, while the dynamics around the baseline remains invariant with respect to the baseline value. Therefore, the network dynamics studied here is to be understood as the dynamics around an arbitrary baseline activity (here set to $r_i = 0$ for simplicity). The term $J_{ij}$ represents the connectivity weight between the presynaptic unit $j$ and the postsynaptic unit $i$. We model left and right trial types through modulations of the external input to the excitatory units, $I_{ext,i}$ assumed to be constant in time. A left trial type corresponds to high input on the left excitatory neuron $E_L$ and no input on the right excitatory neuron $E_R$, and symmetrically for right trials. The term $\eta_i(t)$ represents a source of zero-mean Gaussian input noise to unit $i$, which in general depends on time.

##### The connectivity matrix

For each pre and postsynaptic types $X, Y \in E, I$, we denote by $w_{YX}^{in}$ and $w_{YX}^{out}$ the synapses that connect units belonging to the same or to different subnetworks respectively, and we assume that these connections are symmetric with respect to the left and right subnetworks $L$ and $R$:

$$w_{Y_n X_m}^{in/out} = w_{Y_m X_n}^{in/out}, \; n, m \in L, R \tag{2}$$

This results in a $4 \times 4$ blockwise symmetric connectivity matrix of the form:

$$\mathbf{J} = \begin{pmatrix} \mathbf{W}^{in} & \mathbf{W}^{out} \\ \mathbf{W}^{out} & \mathbf{W}^{in} \end{pmatrix} \tag{3}$$

where the $2 \times 2$ matrices $\mathbf{W}^{in}$ and $\mathbf{W}^{out}$ specify the connectivity weights respectively within a trial-type selective subnetwork and across subnetworks, and are given by:

$$\mathbf{W}^{in} = \begin{pmatrix} w_{EE}^{in} & -w_{EI}^{in} \\ w_{IE}^{in} & -w_{II}^{in} \end{pmatrix}, \; \mathbf{W}^{out} = \begin{pmatrix} w_{EE}^{out} & -w_{EI}^{out} \\ w_{IE}^{out} & -w_{II}^{out} \end{pmatrix}. \tag{4}$$

##### Linear stability analysis

We studied analytically the linear stability of the dynamics of the network with connectivity given by Eq.[3]. The network is stable when all the eigenvalues of the connectivity matrix $\mathbf{J}$ have real part smaller than unity. To compute the eigenvalues $\lambda$ of the connectivity matrix, we solved the eigenvalue equation $\det(\mathbf{J} - \lambda\mathbb{I}) = 0$ for $\lambda$. To simplify calculations, we note that $\mathbf{J} - \lambda\mathbb{I}$ is a blockwise symmetric matrix, so that its determinant can be written as:

$$det(\mathbf{J} - \lambda\mathbb{I}) = \det((\mathbf{W}^{in} - \lambda\mathbb{I}) + \mathbf{W}^{out}) \det((\mathbf{W}^{in} - \lambda\mathbb{I}) - \mathbf{W}^{out}) \tag{5}$$

so that solving $\det(\mathbf{J} - \lambda\mathbb{I}) = 0$ amounts to solving separately the equations:

$$\begin{aligned} \det(\mathbf{W}^{in} + \mathbf{W}^{out} - \lambda\mathbb{I}) = \det(\mathbf{S} - \lambda\mathbb{I}) = 0 \\ \det(\mathbf{W}^{in} - \mathbf{W}^{out} - \lambda\mathbb{I}) = \det(\mathbf{\Delta} - \lambda\mathbb{I}) = 0, \end{aligned} \tag{6}$$

where we defined the matrices $\mathbf{S} = \mathbf{W}^{in} + \mathbf{W}^{out}$ and $\mathbf{\Delta} = \mathbf{W}^{in} - \mathbf{W}^{out}$.
Thus, the four eigenvalues of $\mathbf{J}$ are given by the joint set of the two eigenvalues of $\mathbf{W}^{in} + \mathbf{W}^{out}$ and the two eigenvalues of $\mathbf{W}^{in} - \mathbf{W}^{out}$. Correspondingly, the eigevalues of $\mathbf{J}$ are stable if and only if the eigenvalues of $\mathbf{W}^{in} + \mathbf{W}^{out}$ and those of $\mathbf{W}^{in} - \mathbf{W}^{out}$ are stable. For a general $2 \times 2$ matrix $\mathbf{A}$, its eigenvalues are given by the equation:

$$\lambda_{1,2} = \frac{\mathrm{Tr}(\mathbf{A}) \pm \sqrt{\mathrm{Tr}(\mathbf{A})^2 - 4\det\mathbf{A}}}{2}, \tag{7}$$

while the conditions for their stability are given by:

$$\begin{aligned} \mathrm{Tr}\mathbf{A} < 0 \\ \det\mathbf{A} > 0. \end{aligned} \tag{8}$$

We used Eq.[8] with $\mathbf{A} = \mathbf{W}^{in} \pm \mathbf{W}^{out}$ to derive four necessary and sufficient conditions for the stability of the connectivity matrix $\mathbf{J}$, which read:

$$\begin{aligned} & S_{EE} - S_{II} < 2 \\ & (1 - S_{EE})(1 + S_{II}) + S_{EI}S_{IE} > 0 \\ & \Delta_{EE} - \Delta_{II} < 2 \\ & (1 - \Delta_{EE})(1 + \Delta_{II}) + \Delta_{EI}\Delta_{IE} > 0 \end{aligned} \tag{9}$$

where, for each pre and postsynaptic types $X$ and $Y$, we define

$$\begin{aligned} S_{YX} = w_{YX}^{in} + w_{YX}^{out} \\ \Delta_{YX} = w_{YX}^{in} - w_{YX}^{out}. \end{aligned} \tag{10}$$

The first term, $S_{YX}$, represents twice the average connection strength for within and across-subnetworks connections. The second term, $\Delta_{YX}$, represents the difference in connection strength between within and across-subnetworks connections, which we take as a measure of the selectivity of $X \to Y$ connections: it takes positive values when a specific connection motif is stronger within a subnetwork than across subnetworks ($\Delta_{YX} > 0$: co-selective connection motif), while it takes negative values when the connection motif is stronger across than within a subnetwork ($\Delta_{YX} < 0$: anti-selective connection motif). As explained below, we focused on the connection selectivity $\Delta_{XY}$, as those are the quantities measured in experiments and primarily influence the network dynamics and computations.

#### Parameter exploration

We are interested in how the network dynamics change as a function of the connection selectivity $\Delta_{XY}$. In the next section, we show how the encoding properties of the network depend primarily on the connection selectivities $\Delta_{XY}$. In our analyses we therefore examine the network dynamics and computations as a function of $\Delta_{XY}$ for a fixed $S_{XY}$. Since the terms $w_{YX}^{in/out}$ in Eq.[4] need to be positive and

$$\begin{aligned} w_{XY}^{in} = (S_{XY} + \Delta_{XY})/2 \\ w_{XY}^{out} = (S_{XY} - \Delta_{XY})/2, \end{aligned} \tag{11}$$

the terms $\Delta_{YX}$ must satisfy

$$\Delta_{XY} \in [-S_{XY}, S_{XY}], \tag{12}$$

i.e. they are not constrained in their sign, but are constrained in their magnitude by the terms $S_{XY}$. In Fig. 4b,c,d, Ext. Data Fig. 4 and Supp. Fig 4 and 5, we set $S_{IE} = S_{EI} = 2$, so that $\Delta_{IE} \in [0,2]$ and $\Delta_{EI} \in [-2,2]$. Note that for the E-to-I connections we only consider the region corresponding to positive selectivity. In fact, in absence of inputs on the inhibitory units, the selectivity of these units is inherited by the corresponding excitatory units through E-to-I connections, and the region corresponding to $\Delta_{IE} \in [-2,0]$ leads to the inhibitory unit $I_L$ being selective for the right trial and $I_R$ for the left trial.

#### Response of the network to external inputs

We examined the response of the network to external inputs. We define the network response as the steady-state of the network with external input $\mathbf{I}_{ext}$ and zero noise ($\eta(t) = 0$; or equivalently, as the trial-averaged steady-state response across noisy trials, when zero-mean input noise is present). When the dynamics is linear, the steady-state $r^*$ can be written as a function of the connectivity matrix $\mathbf{J}$ and of the external input $\mathbf{I}_{ext}$ as:

$$r^* = (\mathbb{I} - \mathbf{J})^{-1}\, \mathbf{I}_{ext}. \tag{13}$$

For a connectivity matrix of the form given by Eq.[3] we can compute analytically the steady-state response.

#### Computing the matrix $(\mathbb{I} - \mathbf{J})^{-1}$

We compute the matrix $(\mathbb{I} - \mathbf{J})^{-1}$ by following the method developed in De Mazancout and Gerlic[84] for finding the inverse of block-circulant matrices. Here we follow their notation and apply the method to a $2 \times 2$ block-circulant matrix of the form

$$\mathbf{A} = \begin{pmatrix} \mathbf{a_0} & \mathbf{a_1} \\ \mathbf{a_1} & \mathbf{a_0} \end{pmatrix} \tag{14}$$

where the matrix $\mathbf{A}$ can be mapped in matrix $(\mathbb{I} - \mathbf{J})$ using $\mathbf{a_0} = \mathbb{I} - \mathbf{W}^{in}$ and $\mathbf{a_1} = -\mathbf{W}^{out}$.

The first step of the method involves block-diagonalizing a $2 \times 2$ block matrix $\mathbf{M}$ given by:

$$\mathbf{M} = \begin{pmatrix} \mathbf{0} & \mathbb{I}_2 \\ \mathbb{I}_2 & \mathbf{0} \end{pmatrix}, \tag{15}$$

where $\mathbb{I}_2$ denotes the $2 \times 2$ identity matrix. The block-eigenvalues $\omega_{0,1}$ and the block-eigenvectors $\mathbf{E}_{0,1}$ of $\mathbf{M}$ are given respectively by:

$$\omega_{0,1} = \pm 1,\ \mathbf{E}_{0,1} = (\mathbb{I}_2, \pm\mathbb{I}_2). \tag{16}$$

The second step involves building a matrix $\mathbf{P}$ than contains the block-eigenvectors $\mathbf{E}_{0,1}$ s columns and a matrix $\mathbf{D}$ that contains the block-eigenvalues on the diagonal as:

$$\mathbf{P} = \begin{pmatrix} \mathbb{I}_2 & \mathbb{I}_2 \\ \mathbb{I}_2 & -\mathbb{I}_2 \end{pmatrix},\ \mathbf{D} = \begin{pmatrix} \mathbb{I}_2 & \mathbf{0} \\ \mathbf{0} & -\mathbb{I}_2 \end{pmatrix}. \tag{17}$$

By noting that $\mathbf{P}^{-1} = \mathbf{P}^T/2$, the inverse of the matrix $\mathbf{A}$ can then be written as:

$$\begin{aligned} \mathbf{A}^{-1} &= \mathbf{P}(\mathbf{a_0}\mathbf{D}^0 + \mathbf{a_1}\mathbf{D}^1)^{-1}\mathbf{P}^{-1} \\ &= \mathbf{P}[\mathbf{a_0}\begin{pmatrix} \mathbb{I}_2 & \mathbf{0} \\ \mathbf{0} & \mathbb{I}_2 \end{pmatrix} + \mathbf{a_1}\begin{pmatrix} \mathbb{I}_2 & \mathbf{0} \\ \mathbf{0} & -\mathbb{I}_2 \end{pmatrix}]^{-1}\mathbf{P}^{-1} = \mathbf{P}\begin{pmatrix} (\mathbf{a_0} + \mathbf{a_1})^{-1} & \mathbf{0} \\ \mathbf{0} & (\mathbf{a_0} - \mathbf{a_1})^{-1} \end{pmatrix}\mathbf{P}^{-1} \end{aligned} \tag{18}$$

Thus, we can write:

$$(\mathbb{I}-\mathbf{J})^{-1}=\frac{1}{2}\begin{pmatrix}(\mathbb{I}-\mathbf{S})^{-1}+(\mathbb{I}-\boldsymbol{\Delta})^{-1} & (\mathbb{I}-\mathbf{S})^{-1}-(\mathbb{I}-\boldsymbol{\Delta})^{-1}\\ (\mathbb{I}-\mathbf{S})^{-1}-(\mathbb{I}-\boldsymbol{\Delta})^{-1} & (\mathbb{I}-\mathbf{S})^{-1}+(\mathbb{I}-\boldsymbol{\Delta})^{-1}\end{pmatrix}, \tag{19}$$

where the matrices $\mathbf{S}$ and $\boldsymbol{\Delta}$ are defined by Eq.[6] and

$$\begin{aligned}(\mathbb{I}-\mathbf{S})^{-1}&=\frac{1}{\alpha}\begin{pmatrix}1+S_{II} & -S_{EI}\\ S_{IE} & 1-S_{EE}\end{pmatrix},\ \alpha=(1-S_{EE})(1+S_{II})+S_{EI}S_{IE}\\ (\mathbb{I}-\boldsymbol{\Delta})^{-1}&=\frac{1}{\delta}\begin{pmatrix}1+\Delta_{II} & -\Delta_{EI}\\ \Delta_{IE} & 1-\Delta_{EE}\end{pmatrix},\ \delta=(1-\Delta_{EE})(1+\Delta_{II})+\Delta_{EI}\Delta_{IE},\end{aligned} \tag{20}$$

with the terms $S_{XY}$ and $\Delta_{XY}$ defined by Eq.[10].

When not specified otherwise, in our analysis we focus on the response of excitatory units.

#### Response of the excitatory neurons to external inputs

Using Eq.[19], we can compute the response of the excitatory units $E_L$ and $E_R$ to an external input on both excitatory units ($\mathbf{I}_{ext}=\left(c_{E_L},0,c_{E_R},0\right)$). The response of the excitatory units $r^*=\left(r^*_{E_L},r^*_{E_R}\right)$ is given by:

$$\begin{pmatrix}r^*_{E_L}\\ r^*_{E_R}\end{pmatrix}=\frac{c_{E_L}}{2}\begin{pmatrix}\dfrac{1+S_{II}}{\alpha}+\dfrac{1+\Delta_{II}}{\delta}\\ \dfrac{1+S_{II}}{\alpha}-\dfrac{1+\Delta_{II}}{\delta}\end{pmatrix}+\frac{c_{E_R}}{2}\begin{pmatrix}\dfrac{1+S_{II}}{\alpha}-\dfrac{1+\Delta_{II}}{\delta}\\ \dfrac{1+S_{II}}{\alpha}+\dfrac{1+\Delta_{II}}{\delta}\end{pmatrix} \tag{21}$$

#### Amplification through opponent inhibition

In this section we examine how the amplification of trial-type signal depends on the degree of opponent inhibition between excitatory units.

We model different trial types by differentially modulating the external inputs to the excitatory units. Left (resp. right) trials correspond to high input on the left (resp. right) excitatory unit and low input to the right (resp. left) unit, i.e. $\mathbf{I}_{ext}=\left(c_{E_L},0,c_{E_R},0\right)$ with $c_{E_L}>c_{E_R}$ for left trials, while $c_{E_R}>c_{E_L}$ for right trials. For simplicity, and without loss of generality, we assume that $c_{E_L}=1, c_{E_R}=0$ (resp. $c_{E_L}=0, c_{E_R}=1$) for left (resp. right) trial types. In addition, in this analysis we neglect E-to-E and I-to-I connectivity to examine the contributions of the E-to-I and I-to-E connections to the network response. The overall results hold when including E-to-E and I-to-I connections (Supp. Fig. 4).

Using Eq.[21] we can write the response of the network during left trial types as:

$$\begin{pmatrix}r^*_{E_L}\\ r^*_{E_R}\end{pmatrix}=\frac{1}{2}\begin{pmatrix}1/\alpha+1/\delta\\ 1/\alpha-1/\delta\end{pmatrix}=\frac{1}{2}\begin{pmatrix}\alpha^{-1}+(1+\Delta_{EI}\Delta_{IE})^{-1}\\ \alpha^{-1}-(1+\Delta_{EI}\Delta_{IE})^{-1}\end{pmatrix} \tag{22}$$

where we made explicit the terms that depend on the selectivity of E-to-I and I-to-E connections, $\Delta_{IE}$ and $\Delta_{EI}$. To quantify the contribution of different connection selectivities to the network response, we fix the average connection weights by fixing the quantities $S_{XY}$, and vary systematically the connection selectivities $\Delta_{YX}$ (see Method Section "Parameter exploration").

We note that the second term of the sums in Eq.[22] increases as $\Delta_{EI}\Delta_{IE}$ decreases. In particular, when $\Delta_{EI}\Delta_{IE}<0$, the term $1/\delta$ is larger than one, and becomes very large for $\Delta_{EI}\Delta_{IE}\gtrsim-1$ (where $\Delta_{EI}\Delta_{IE}=-1$ corresponds to the stablity boundary). When $1/\delta$ is large ($\delta\ll\alpha$), an input on the left selective excitatory unit $E_L$ has the effect of suppressing the unit with opposite trial-type selectivity $E_R$, which in turn enhances the activity of the left selective neuron $E_L$. This mechanism is mediated by opponent inhibition between the two trial-selective excitatory units, and is enhanced when the cross-inhibition between these two units is stronger than the inhibition acting within one single pool. To illustrate this point, we computed the difference of the inhibition acting within one single pool $\text{Inh}_{in}$ and the inhibition acting across pools $\text{Inh}_{out}$, as the sum of single contributions from distinct second-order inhibitory connectivity motifs, and obtained:

$$\begin{aligned} \text{Inh}_{in} - \text{Inh}_{out} &= \left[w_{IE}^{in} w_{EI}^{in} + w_{IE}^{out} w_{EI}^{out}\right] - \left[w_{IE}^{in} w_{EI}^{out} + w_{IE}^{out} w_{EI}^{in}\right] \\ &= \Delta_{EI}\Delta_{IE} \end{aligned} \tag{23}$$

Thus, the term $\Delta_{EI}\Delta_{IE}$ appearing in Eq.[22] corresponds to first approximation to the difference between the inhibition within and the inhibition across pools. Negative values of $\Delta_{EI}\Delta_{IE}$ therefore indicate that, to the lowest order in the connectivity motifs, the across-pool inhibition is stronger than the inhibition acting within a pool.

We focus our analysis on co-selective E-to-I motifs, for which the inhibitory neuron $I_L$ (resp. $I_R$) are selective for the left (resp. right) trials. Thus, for fixed co-selective E-to-I connections, increasing the anti-selectivity of I-to-E motifs increases the suppression of the opposite selective unit as well as the amplification of the unit receiving trial-specific input through opponent inhibition between the two units. Thus, this opponent inhibition determines a disinhibitory motif that effectively acts as a positive feedback within each excitatory pool, amplifying selective external input received by the excitatory units.

#### Encoding of trial-type signal in the network activity

In this section, we describe how the trial-type signal is encoded in the population activity of the excitatory units. In particular, we examine the dimension of the neural state space along which the separation of trial-specific activity occurs, which corresponds to the dimension encoding the trial-type signal.

We assume that the external trial-specific inputs are symmetric for left and right trials, i.e. for left trials $\mathbf{I}_L = (c_1, 0, c_2, 0)$ with $c_1 > c_2$, while for right trials the inputs on the excitatory neurons are reversed, i.e. $\mathbf{I}_R = (c_2, 0, c_1, 0)$, with $c_1 > c_2$. Under this scenario, we can compute the trial-type encoding dimension, defined as the vector connecting the trial-specific mean responses of the excitatory neurons on left and right trials. In particular, we compute the encoding dimension at the input and output stage defined by:

$$\begin{aligned} \mathbf{d}_{in} &= \mathbf{I}_L - \mathbf{I}_R \\ \mathbf{d}_{out} &= \mathbf{r}_L^* - \mathbf{r}_R^*, \end{aligned} \tag{24}$$

where $\mathbf{r}_{L/R}^*$ are the responses of the network to inputs $\mathbf{I}_{L/R}$. The input encoding dimension can be understood as the dimension connecting trial-specific responses in the absence of recurrent connections, in which case the response of the excitatory neurons would equal their external inputs.

We can easily find the input encoding dimension restricted to the excitatory population $(E_L, E_R)$ as:

$$\mathbf{d}_{in} = (c_1 - c_2, -c_1 + c_2). \tag{25}$$

Using Eq.[21] we obtain the output encoding dimension as:

$$\begin{aligned} \mathbf{d}_{out} &= \frac{1}{2}\begin{bmatrix} c_1(\alpha^{-1} + \delta^{-1}) + c_2(\alpha^{-1} - \delta^{-1}) \\ c_1(\alpha^{-1} - \delta^{-1}) + c_2(\alpha^{-1} + \delta^{-1}) \end{bmatrix} - \frac{1}{2}\begin{bmatrix} c_2(\alpha^{-1} + \delta^{-1}) + c_1(\alpha^{-1} - \delta^{-1}) \\ c_2(\alpha^{-1} - \delta^{-1}) + c_1(\alpha^{-1} + \delta^{-1}) \end{bmatrix} \\ &= \frac{1}{\delta}\begin{bmatrix} c_1 - c_2 \\ -c_1 + c_2 \end{bmatrix}, \end{aligned} \tag{26}$$

where $\delta = (1 - \Delta_{EE})(1 + \Delta_{II}) + \Delta_{EI}\Delta_{IE}$ (see Eq.[20]). From Eq.[25] and Eq.[26], we obtain the relationship between the input and output encoding dimensions, given by:

$$\mathbf{d}_{out} = \frac{1}{\delta}\,\mathbf{d}_{in}. \tag{27}$$

Eq.[27] reveals how the vector that encodes the trial type signal changes from input to output due to the recurrent network connectivity. First, we note that the network dynamics do not change the direction of the encoding vector, i.e. the encoding vector maintains the same direction set by the external inputs. Second, the recurrent connectivity modulates the separation (or distance) between the mean activities according to $||\mathbf{d}_{out}||/||\mathbf{d}_{in}|| = 1/\delta$. On one hand, when $\delta$ decreases the separation between output mean activities $\mathbf{d}_{out}$ increases. On the other hand, when $\delta < 1$ the separation between trial-specific activities is larger at the output than at the input stage. Importantly, under the assumption of symmetric inputs for left and right trial types, the modulation of the encoding vector through the recurrent synapses only depends on the selectivity of the connection $\Delta_{XY}$ and not on the average magnitude of the connection represented by $S_{XY}$.
In the absence of E-to-E and I-to-I connections, increased separation of trial-specific activities occurs when $\Delta_{EI}\Delta_{IE} < 0$, corresponding to co-selective E-to-I and anti-selective I-to-E.

#### Robustness of the trial-type encoding to noise

In the previous sections we have shown that the opponent inhibition implemented by oppositely-selective inhibitory motifs has the effect of increasing the separation of the trial-specific mean activities with respect to equally selective motifs on one hand, and of increasing the output separation with respect to the input separation on the other hand (Eq.[27], Ext. Data Fig. 4a-c). In this section we study how this impact the robustness of the trial-type encoding to different sources of noise. Specifically, we will focus on two types of noise, i.e., readout noise, acting at the level of a hypothetical readout unit that optimally decodes the trial-type signal on single trials, and input noise, acting at the level of the input to the excitatory units. The effect of different connectivity motifs on the noise robustness is quantified by computing the decoding accuracy of a readout unit trained to optimally classify the trial type from the neural activity of the excitatory units on single noisy trials, since excitatory units are those that project to downstream areas. We assume that the noise has zero mean and Gaussian statistics.

#### Decoding accuracy

We analytically compute the performance of an optimal linear decoder trained to classify trial types from the neural activity of the excitatory units as follows. Given the difference in mean activity between left and right trial types $\Delta\boldsymbol{\mu}$, and the noise covariance matrix $\mathbf{C}$ (which for linear systems does not depend on the identity of the trial type), we compute the signal-to-noise ratio as:

$$SNR = \Delta\boldsymbol{\mu}^T\mathbf{C}^{-1}\Delta\boldsymbol{\mu}. \tag{28}$$

The decoding accuracy of an optimal linear classifier can be computed as a function of the signal-to-noise ratio $SNR$ as:

$$\text{Decod. acc.} = \Phi\left(\sqrt{\frac{SNR}{4}}\right), \text{ where } \Phi(x) = \int_{-\infty}^{x} \frac{e^{z^2/2}}{\sqrt{2\pi\sigma^2}} dz. \tag{29}$$

In Fig. 4c,d and Supp. Fig. 5 we defined the relative decoding accuracy as the ratio of the output decoding accuracy computed using the network steady-state activity and the input decoding accuracy computed using the inputs to the network, after subtracting the chance decoding accuracy:

$$Relative\ decod.acc. = \frac{Decod.acc.(output) - 0.5}{Decod.acc.(input) - 0.5}.$$

#### Noise in the readout

In this section, we investigate the effects of single-trial variability on decoding accuracy when decoding the trial type from single-trial responses. We thus trained an optimal linear readout neuron to decode the trial type from the single-trial responses of the excitatory units. Importantly, we assume that the trial-to-trial variability only affects the readout activity, so that the noise covariance matrix is fixed across the range of network parameters. If the optimal decoder weights are given by the vector $\mathbf{w} = \left(w_{E_L}, w_{E_R}\right)$, and the neural activity on a trial $\nu$ is given by $\mathbf{r}$, this trial will be classified into a specific trial-type class according to the activity of the linear readout $z$ given by:

$$z = \sum_{i=E_L,E_R} w_i\, r_i + \sigma_z\, \xi,\ with\ \langle \xi^2 \rangle = 1 \tag{30}$$

where the second term on the right-hand side represents the readout noise. Trial $\nu$ is then classified as left or right trial type if $z > \theta$ or $z < \theta$, where $\theta$ is a bias term. Effectively, we model the readout noise by generating the mean responses for each trial type, then adding uncorrelated Gaussian noise on the mean responses to generate single-trial responses, which are classified by the linear decoder. Using this procedure, the readout activity can be written as

$$z = \sum_{i=E_L,E_R} w_i\,(r_i + \sigma_r\, \xi_i),\ \langle \xi_i \xi_j \rangle = \delta_{ij}. \tag{31}$$

Note that this procedure is equivalent to adding noise directly to the readout activity $z$. In fact, Eq.[30] and Eq.[31] coincide if $\sigma_z^2 = \sigma_r^2\left(w_{E_L}^2 + w_{E_R}^2\right)$.

If the trial-to-trial variability is fixed across the range of network parameters, an increase in the mean response separation due to increased opponent inhibition between selective units, always results in a higher decoding performance, since higher separation of mean activities increases the linear separability of the single-trial responses on different trial types, therefore enhancing the decoding accuracy. In fact, from Eq.[31] it follows that $\mathbf{C}^{-1} = \sigma_r^{-2}\mathbb{I}$, so that $SNR = \sigma_r^{-2}||\Delta\boldsymbol{\mu}||^2$ and the decoding accuracy is given by

$$\text{Decod. acc.} = \Phi\left(\frac{||\Delta\boldsymbol{\mu}||}{2\sigma_r}\right), \tag{32}$$

which is a monotonically increasing function of the distance between the mean activities $||\Delta\boldsymbol{\mu}||$.
We measured the decoding accuracy computed as in Eq.[29] as a function of the connection selectivity $\Delta_{XY}$. Results show that regions of the parameter space where the distance between the mean activities is larger correspond to higher absolute decoding accuracy (Fig. 4c,d and Ext. Data Fig. 4c,d,e).

We then quantified to which extent specific patterns of connection selectivity $\Delta_{XY}$ are beneficial for separating input signals. We computed the ratio between the decoding accuracy evaluated for output responses and the decoding accuracy evaluated for input patterns (see Method Section "Decoding accuracy"). Values larger than unity indicate that the recurrent connections were able to increase the separation of the input patterns, therefore increasing linear decodability. The condition for the relative decoding accuracy to be larger than unity can be obtained using Eq.[27]. Since for readout noise the decoding accuracy is a monotonic function of the separation between mean activities, we obtain that the relative decoding accuracy is larger than one if the relative distance $\mathbf{d}_{out}/\mathbf{d}_{in}$ is larger than one. This happens in the regions of parameters where $\delta < 1$ (while $\delta > 0$ for stability). In absence of E-to-E and I-to-I connections, this condition corresponds to $\Delta_{EI}\Delta_{IE} < 0$, i.e. to co-selective E-to-I and anti-selective I-to-E motifs.

#### Noise in the input signal

Here we examine how noise in the input signal affects the encoding of the trial type for different values of connection selectivity. In general, the input signal may exhibit trial-to-trial variability and, in networks that amplify input signals, signal amplification may be accompanied by amplification of this variability. Examining the interplay between signal and noise amplification is therefore crucial to characterize the robustness of signal amplification to noise in the input. We consider two distinct types of noise $\eta(t)$: stationary and non-stationary noise.

#### Non-stationary noise

We consider Gaussian non-stationary noise with zero mean and uncorrelated variability across trials $\nu$'s and across time $t$:

$$\eta(t) = \sigma\, \xi^{\nu}(t),\ \langle \xi^{\nu_1}(t)\xi^{\nu_2}(t')\rangle = \delta_{\nu_1\nu_2}\delta(t-t'). \tag{33}$$

For each combination of network parameters, we computed the decoding accuracy using Eq.[28] and [29]. The vectors defininf the differences in mean activities on right and left trials at the input and output stages are given by

$$\begin{aligned} \Delta\boldsymbol{\mu}_{in} &= \mathbf{I}_{ext} \\ \Delta\boldsymbol{\mu}_{out} &= \mathbf{M}\,\Delta\boldsymbol{\mu}_{in},\ \boldsymbol{with}\ \mathbf{M} = (\mathbb{I}-\mathbf{J})^{-1}. \end{aligned} \tag{34}$$

We computed the covariance matrix $\mathbf{C}_{out}$ of the network response at large times by numerically solving the Lyapunov equation

$$\mathbf{M}\mathbf{C}_{out} + \mathbf{C}_{out}\mathbf{M}^T = \sigma^2\mathbb{I}. \tag{35}$$

We found that in the parameter regions corresponding to opponent inhibition, both absolute and relative decoding accuracy were enhanced (Supp. Fig. 5). This is explained by the fact that, for non-stationary noise, both the signal and the noise are amplified by the recurrent connections, but the signal amplification is larger than the noise amplification. This can be illustrated by a toy model of a single neuron with an excitatory autapse with weight equal to $w$. The response of the network to an input $I$ is given by $r^* = I_{ext}/(1-w)$, while its variance is given by $Var(r^*) = \sigma^2/[2(1-w)]$. Thus, the signal and the variance increase in the same way ($\sim 1/(1-w)$), so that the signal-to-noise ratio increases with the excitatory weight $w$ as:

$$SNR \sim r^*/\sqrt{var(r^*)} \sim 1/\sqrt{1-w}. \tag{36}$$

The same principle holds for a two-units toy model that has opponent inhibition between the units.

As a result, for non-stationary input noise, the recurrent connectivity can amplify the signal more than it amplifies the noise. Networks that strongly amplify the trial-type signal are therefore robust to non-stationary input noise and can exhibit enhanced absolute and relative decoding accuracy for trial-type classification.

#### Stationary noise

A second scenario corresponds to noise that does not vary in time and is therefore stationary. In this case, for a specific trial $\nu$, the external input to the system will have a noise contribution given by:

$$\eta(t) = \sigma\xi^{\nu},\ \langle \xi^{\nu_1}\xi^{\nu_2}\rangle = \delta_{\nu_1\nu_2}, \tag{37}$$

where the noise has zero mean and is uncorrelated across trials. Here we show that, in a linear network with stationary noise, the information present in the input is equal to the information present at the output stage. This is a consequence of linear dynamics and of the fact that stationary variations of the inputs are amplified to the same extent as the mean input, leading to unchanged information content at the input and at the output. As a result, the information at the output is the same for every combination of the network parameters, and equal to the input information. To show this, let $\Delta\boldsymbol{\mu}_{in}$ and $\Delta\boldsymbol{\mu}_{out}$ be the mean activity separation at the input and at the output, and let $\mathbf{C}_{in}$ = and $\mathbf{C}_{out}$ be the covariance matrices of the input and output responses across trials. Since at steady-state the response of the network is given by

$$\mathbf{r}^* = \mathbf{M}(\mathbf{I}_{ext} + \sigma\boldsymbol{\xi}),\ \mathbf{M} = (\mathbb{I} - \mathbf{J})^{-1} \tag{38}$$

we have that the mean input and output activities are given by Eq.[34] and the input and output covariance matrices can be written as

$$\begin{aligned} \mathbf{C}_{in} &= \sigma^2\mathbb{I} \\ \mathbf{C}_{out} &= \mathbf{M}\mathbf{C}_{in}\mathbf{M}^T. \end{aligned} \tag{39}$$

Using these relationships, we can compute the signal-to-noise ratio at the input and output stage as

$$\begin{aligned} SNR_{in} &= \mathbf{I}_{ext}^T\mathbf{C}_{in}^{-1}\mathbf{I}_{\mathbf{ext}} \\ SNR_{out} &= \Delta\boldsymbol{\mu}_{out}^T\mathbf{C}_{out}^{-1}\Delta\boldsymbol{\mu}_{out} = \mathbf{I}_{ext}^T\mathbf{M}^T(\mathbf{M}\mathbf{C}_{in}\mathbf{M}^T)^{-1}\mathbf{M}\mathbf{I}_{ext} = \mathbf{I}_{ext}^T\mathbf{C}_{in}^{-1}\mathbf{I}_{ext} = SNR_{in} \end{aligned} \tag{40}$$

Therefore, when decoding the trial type from the full neural population (excitatory and inhibitory neurons), the resulting decoding accuracy at the output is equal to the decoding accuracy at the input regardless of the network parameters, and thus does not change as a function of the network parameters. Even when the decoding accuracy was computed using only the activity of the excitatory population, we observed weak modulation of the accuracy with the network parameters (Supp. Fig. 5). Importantly, we did not observe an enhancement of the decoding accuracy in the region corresponding to opponent inhibition, as we did in the case of readout noise and non-stationary input noise.

Thus, networks that amplify input signal also amplify input noise that is stationary in time. For stationary input noise, signal amplification does not provide an advantage in terms of absolute nor relative decoding accuracy (Supp. Fig. 5). In contrast, networks based on opponent inhibition can enhance decoding accuracy when the trial-to-trial variability is due to noise in the input that varies in time or to noise in the readout activity. In these cases, both the absolute decoding accuracy and the relative decoding accuracy are enhanced in parameter regions corresponding to networks that implement signal amplification through opponent inhibitory motifs shaped by co-selective E-to-I and anti-selective I-to-E connectivity motifs.

#### The recurrent neural network model

To test if the predictions of the linear rate model hold for a more realistic higher-dimensional network model able to generate PPC activity, we trained a recurrent neural network (RNN) model to reproduce the trial-averaged PPC calcium traces on left and right trials. We then analysed the connectivity weights obtained after training to test the predictions of the linear rate model on the trained networks. The training procedure we adopt here is inspired by the work of Rajan and colleagues[53], and we refer to their work for a detailed account of the method. Below we give a concise overview of the RNN setup and related analyses.

##### Network setup

We train an RNN to reproduce the PPC activity traces. The RNN is described by the differential equation

$$\tau\dot{x}_i = -x_i + \sum_{j=1}^{N} J_{ij}\,\phi(x_j) + \sigma\xi_i(t), \tag{41}$$

where $x_i$ represent the synaptic current of neuron $i$ and $r_i = \phi(x_i)$ is the firing rate of neuron $i$, and the last term on the right-hand side is a source of non-stationary Gaussian uncorrelated external noise. The network is constrained to have the same number of neurons and the same type (E or I) as the experimentally reconstructed circuit ($N = 137, N_E = 121, N_I =$

16). Accordingly, the connectivity matrix $J_{ij}$ is constrained to satisfy Dale's law, i.e. with elements belonging to the same column having the same sign according to the type of the pre-synaptic neurons[54].

We assume the nonlinearity $\phi(x)$ to be a softplus function (smoothed ReLU) given by:

$$\phi(x) = \frac{\ln\left(1 + e^{k(x-\theta)}\right)}{k}, \tag{42}$$

with the threshold $\theta = 1$ and the sharpness parameter $k = 0.5$.

Prior to training, the PPC trial-averaged calcium activity traces were normalized to the peak of each cell's activity on preferred trials. We trained several (n=192) network realizations corresponding to different independent realizations of the external inputs $\xi_i(t)$ and different initializations of the connectivity matrix. The connectivity matrix was initialized as a random Gaussian matrix with zero mean and standard deviation equal to $g = 0.1$.

#### External inputs

The external input consisted in filtered white noise generated according to the equation

$$\tau_{WN} \dot{I}_{ext,i} = -I_{ext,i} + a_i \, \eta_i(t), \tag{43}$$

where $\eta(t)$ is a Gaussian white noise with zero mean and unit variance, and $\tau_{WN} = 1$. The input is fed only to the excitatory neurons. Inputs to the inhibitory neurons were set to zero. We modelled different trial types by modulating the amplitude and initial conditions of the input noise on the excitatory neurons. For a left trial, we set $a_i = 1$ and $I_{ext,i} = 1$ if neuron $i$ was a left selective excitatory neuron, or $a_i = 0.25$ and $I_{ext,i} = 0.25$ if neuron $i$ was a right selective one, and symmetrically for right trials.

#### Connection selectivity

For each connection motif, we quantified connection selectivity by computing the Spearman's correlation between the connectivity weights and the selectivity similarity. We computed the selectivity similarity as for the connectomics data, i.e. for each pair of neurons with selectivity indices $X_i$ and $X_j$ the selectivity similarity was computed as

$$\mathrm{ss}_{ij} = sgn(X_i X_j)\sqrt{|X_i||X_j|}. \tag{44}$$

We then compute the Spearman's correlation between the connectivity weights $J_{ij}$ and $ss_{ij}$, which defined the connection selectivity.

#### Perturbation of the connectivity weights

For each connection type $X \to Y$ we perturbed the selectivity $\Delta_{YX}$ by keeping the average connection strength $S_{YX}$ constant, as described below. We continuously varied the perturbation parameters $\delta_{in}$ and $\delta_{out}$ so that within- and across-subnetwork connection weights were re-scaled according to

$$\begin{aligned} w_{YX}^{in} &\longrightarrow (1 + \delta_{in}) \, w_{YX}^{in} \\ w_{YX}^{out} &\longrightarrow (1 + \delta_{out}) \, w_{YX}^{out}. \end{aligned} \tag{45}$$

To ensure that $S_{YX} = w_{YX}^{in} + w_{YX}^{out}$ be constant when varying $\delta_{in}$, the perturbation parameter for the across-subnetwork connections needs to be given by:

$$\delta_{out} = -\frac{\langle w_{YX}^{in} \rangle}{\langle w_{YX}^{out} \rangle} \delta_{in}, \tag{46}$$

where $\left\langle w_{YX}^{in/out} \right\rangle$ represents the average within- and across-subnetwork weight. In our analyses, we varied $\delta_{in}$ in the interval $[-0.1, 0.1]$.

For each value of $\delta_{in}$, we regenerated the network dynamics using the new set of connectivity weights. We then computed the distance between the population neural trajectories on left and right trials, then averaged the distance over time. In Fig 4i we show the value of the distance as a function of the perturbed value of the selectivity $\Delta_{EI}$ or $\Delta_{IE}$ normalized by the value of the distance corresponding to the original trained network.

### Supplemental Tables

#### Supplementary Table 1: E-to-I connections

| source neuron | source selectivity index | target neuron | target selectivity index | selectivity similarity index | number of synapses | cable overlap (μm) | norm. syn. freq. ($\mu m^{-1}$) | avg. PSD area ($\mu m^2$) |
|---|---|---|---|---|---|---|---|---|
| 143494 | -0.0845 | 8357 | 0.112 | -0.0975 | 1 | 90.7 | 0.011 | 0.0378 |
| 144125 | -0.0656 | 8357 | 0.112 | -0.0859 | 2 | 145 | 0.0138 | 0.133 |
| 7287 | -0.0632 | 8357 | 0.112 | -0.0843 | 1 | 21.3 | 0.047 | 0.0532 |
| 143672 | -0.0427 | 140120 | 0.142 | -0.0778 | 2 | 69.1 | 0.0289 | 0.153 |
| 22895 | -0.0763 | 24051 | 0.0479 | -0.0605 | 1 | 69.1 | 0.0145 | 0.101 |
| 173105 | -0.0117 | 140120 | 0.142 | -0.0407 | 1 | 37.1 | 0.0269 | 0.131 |
| 175383 | -0.00228 | 8357 | 0.112 | -0.016 | 1 | 58.9 | 0.017 | 0.325 |
| 144141 | -0.00401 | 24051 | 0.0479 | -0.0139 | 1 | 32.7 | 0.0306 | 0.0818 |
| 168526 | 0.0232 | 175761 | 0.101 | 0.0485 | 1 | 23 | 0.0435 | 0.0482 |
| 171031 | 0.056 | 24051 | 0.0479 | 0.0518 | 1 | 94.3 | 0.0106 | 0.302 |
| 150951 | 0.0603 | 24051 | 0.0479 | 0.0537 | 1 | 21.9 | 0.0456 | 0.229 |
| 144403 | 0.0305 | 140120 | 0.142 | 0.0658 | 1 | 19.8 | 0.0505 | 0.0941 |
| 143985 | 0.0508 | 10607 | 0.0999 | 0.0712 | 1 | 25.4 | 0.0394 | 0.202 |
| 144425 | 0.0466 | 8357 | 0.112 | 0.0724 | 1 | 10.2 | 0.0979 | 0.1 |
| 141671 | 0.0557 | 8357 | 0.112 | 0.0791 | 2 | 84.3 | 0.0237 | 0.21 |
| 197166 | 0.0507 | 140120 | 0.142 | 0.0848 | 1 | 15.4 | 0.065 | 0.14 |
| 143985 | 0.0508 | 140120 | 0.142 | 0.0849 | 1 | 14.8 | 0.0675 | 0.295 |
| 175650 | 0.301 | 24051 | 0.0479 | 0.12 | 4 | 96 | 0.0417 | 0.232 |
| 143896 | 0.263 | 141681 | 0.0895 | 0.153 | 2 | 28.1 | 0.0711 | 0.575 |
| 22901 | 0.167 | 140120 | 0.142 | 0.154 | 2 | 41.1 | 0.0486 | 0.188 |
| 175650 | 0.301 | 10607 | 0.0999 | 0.173 | 1 | 16.8 | 0.0594 | 0.113 |

#### Supplementary Table 2: I-to-E connections

| source neuron | source selectivity index | target neuron | target selectivity index | selectivity similarity index | number of synapses | cable overlap (μm) | norm. syn. freq. ($\mu m^{-1}$) | avg. PSD area ($\mu m^2$) |
|---|---|---|---|---|---|---|---|---|
| 26266 | -0.538 | 143896 | 0.263 | -0.376 | 2 | 77 | 0.026 | 0.127 |
| 26266 | -0.538 | 158983 | 0.0686 | -0.192 | 1 | 105 | 0.00956 | 0.0518 |
| 140120 | 0.142 | 177789 | -0.192 | -0.165 | 2 | 56 | 0.0357 | 0.142 |
| 26266 | -0.538 | 159246 | 0.0382 | -0.143 | 1 | 11.4 | 0.0874 | 0.124 |
| 140120 | 0.142 | 143717 | -0.144 | -0.143 | 2 | 81.3 | 0.0246 | 0.12 |
| 140120 | 0.142 | 144005 | -0.144 | -0.143 | 1 | 32.8 | 0.0305 | 0.0982 |
| 10607 | 0.0999 | 151057 | -0.174 | -0.132 | 1 | 22.4 | 0.0446 | 0.0653 |
| 141681 | 0.0895 | 143717 | -0.144 | -0.114 | 1 | 76.3 | 0.0131 | 0.169 |
| 5141 | 0.12 | 190907 | -0.107 | -0.113 | 2 | 47.8 | 0.0418 | 0.074 |
| 10607 | 0.0999 | 142679 | -0.113 | -0.106 | 1 | 46.2 | 0.0216 | 0.105 |
| 190640 | -0.0402 | 58 | 0.227 | -0.0956 | 1 | 150 | 0.00668 | 0.153 |
| 10607 | 0.0999 | 144461 | -0.0765 | -0.0874 | 1 | 16.8 | 0.0594 | 0.0957 |
| 10607 | 0.0999 | 22895 | -0.0763 | -0.0873 | 1 | 58.6 | 0.0171 | 0.0526 |
| 141681 | 0.0895 | 143785 | -0.0845 | -0.087 | 2 | 77.6 | 0.0258 | 0.102 |
| 24051 | 0.0479 | 143717 | -0.144 | -0.0832 | 2 | 102 | 0.0195 | 0.0952 |
| 10607 | 0.0999 | 190225 | -0.0674 | -0.0821 | 1 | 37.6 | 0.0266 | 0.115 |
| 10607 | 0.0999 | 23919 | -0.0558 | -0.0747 | 1 | 246 | 0.00407 | 0.0492 |
| 175761 | 0.101 | 180507 | -0.0497 | -0.071 | 1 | 95.5 | 0.0105 | 0.16 |
| 190640 | -0.0402 | 25998 | 0.0965 | -0.0623 | 4 | 145 | 0.0276 | 0.255 |
| 24051 | 0.0479 | 22895 | -0.0763 | -0.0605 | 1 | 152 | 0.0066 | 0.105 |
| 24051 | 0.0479 | 190225 | -0.0674 | -0.0568 | 1 | 50.8 | 0.0197 | 0.0888 |
| 24051 | 0.0479 | 144125 | -0.0656 | -0.0561 | 1 | 27.3 | 0.0366 | 0.111 |
| 24051 | 0.0479 | 7287 | -0.0632 | -0.055 | 1 | 77.6 | 0.0129 | 0.0875 |
| 24051 | 0.0479 | 174062 | -0.0591 | -0.0532 | 1 | 18.8 | 0.0531 | 0.0802 |
| 24051 | 0.0479 | 23919 | -0.0558 | -0.0517 | 1 | 87.9 | 0.0114 | 0.103 |
| 24051 | 0.0479 | 190198 | -0.0523 | -0.05 | 3 | 126 | 0.0238 | 0.159 |
| 24051 | 0.0479 | 16384 | -0.051 | -0.0494 | 3 | 52.4 | 0.0573 | 0.163 |
| 190640 | -0.0402 | 171031 | 0.056 | -0.0474 | 1 | 32.6 | 0.0306 | 0.0818 |
| 190640 | -0.0402 | 173088 | 0.0516 | -0.0456 | 1 | 165 | 0.00606 | 0.0694 |
| 190640 | -0.0402 | 143985 | 0.0508 | -0.0452 | 1 | 53.2 | 0.0188 | 0.364 |
| 140120 | 0.142 | 173105 | -0.0117 | -0.0407 | 1 | 97.2 | 0.0103 | 0.412 |
| 5141 | 0.12 | 173105 | -0.0117 | -0.0374 | 1 | 34.8 | 0.0287 | 0.151 |

| | | | | | | | | |
|---|---|---|---|---|---|---|---|---|
| 190640 | -0.0402 | 143250 | 0.031 | -0.0353 | 3 | 82.5 | 0.0364 | 0.177 |
| 141681 | 0.0895 | 173105 | -0.0117 | -0.0323 | 2 | 227 | 0.00882 | 0.224 |
| 24051 | 0.0479 | 173105 | -0.0117 | -0.0237 | 1 | 27.2 | 0.0367 | 0.0806 |
| 24051 | 0.0479 | 159069 | -0.0116 | -0.0235 | 1 | 38.7 | 0.0259 | 0.0693 |
| 144279 | -0.015 | 144403 | 0.0305 | -0.0214 | 1 | 35.7 | 0.028 | 0.456 |
| 24051 | 0.0479 | 144141 | -0.00401 | -0.0139 | 1 | 73.9 | 0.0135 | 0.299 |
| 190640 | -0.0402 | 143483 | 0.00213 | -0.00926 | 4 | 87.3 | 0.0458 | 0.14 |
| 144279 | -0.015 | 144452 | 1.52E-05 | -0.000478 | 1 | 46.1 | 0.0217 | 0.291 |
| 24051 | 0.0479 | 176218 | 0.00102 | 0.007 | 1 | 13.1 | 0.0763 | 0.0716 |
| 190640 | -0.0402 | 175383 | -0.00228 | 0.00958 | 1 | 123 | 0.00813 | 0.137 |
| 5141 | 0.12 | 176218 | 0.00102 | 0.0111 | 1 | 23.4 | 0.0427 | 0.155 |
| 190640 | -0.0402 | 144141 | -0.00401 | 0.0127 | 2 | 85.3 | 0.0234 | 0.206 |
| 141681 | 0.0895 | 158745 | 0.00359 | 0.0179 | 2 | 180 | 0.0111 | 0.107 |
| 5141 | 0.12 | 158745 | 0.00359 | 0.0207 | 1 | 57.6 | 0.0174 | 0.173 |
| 190640 | -0.0402 | 173105 | -0.0117 | 0.0217 | 1 | 65.5 | 0.0153 | 0.169 |
| 24051 | 0.0479 | 4852 | 0.012 | 0.024 | 1 | 33.6 | 0.0297 | 0.172 |
| 190640 | -0.0402 | 147 | -0.0264 | 0.0326 | 3 | 237 | 0.0126 | 0.122 |
| 24051 | 0.0479 | 143250 | 0.031 | 0.0385 | 2 | 257 | 0.00777 | 0.139 |
| 10607 | 0.0999 | 7506 | 0.0163 | 0.0403 | 1 | 21.2 | 0.0471 | 0.235 |
| 190640 | -0.0402 | 143672 | -0.0427 | 0.0414 | 1 | 96.2 | 0.0104 | 0.18 |
| 190640 | -0.0402 | 198820 | -0.0442 | 0.0421 | 1 | 13.5 | 0.0743 | 0.0453 |
| 190640 | -0.0402 | 16384 | -0.051 | 0.0453 | 2 | 131 | 0.0153 | 0.199 |
| 24051 | 0.0479 | 144366 | 0.0443 | 0.0461 | 1 | 105 | 0.0095 | 0.0784 |
| 26266 | -0.538 | 144141 | -0.00401 | 0.0464 | 1 | 11.7 | 0.0851 | 0.0577 |
| 190640 | -0.0402 | 174062 | -0.0591 | 0.0487 | 1 | 105 | 0.00953 | 0.0668 |
| 190640 | -0.0402 | 143810 | -0.0591 | 0.0488 | 3 | 222 | 0.0135 | 0.21 |
| 24051 | 0.0479 | 143985 | 0.0508 | 0.0493 | 2 | 72.8 | 0.0275 | 0.093 |
| 190640 | -0.0402 | 144125 | -0.0656 | 0.0514 | 1 | 59.6 | 0.0168 | 0.149 |
| 24051 | 0.0479 | 171031 | 0.056 | 0.0518 | 1 | 38.5 | 0.0259 | 0.106 |
| 190640 | -0.0402 | 175865 | -0.0678 | 0.0522 | 1 | 60.8 | 0.0164 | 0.415 |
| 141681 | 0.0895 | 144403 | 0.0305 | 0.0523 | 1 | 69.4 | 0.0144 | 0.444 |
| 24051 | 0.0479 | 5150 | 0.0603 | 0.0537 | 2 | 120 | 0.0167 | 0.151 |
| 24051 | 0.0479 | 159960 | 0.0651 | 0.0558 | 1 | 95.6 | 0.0105 | 0.139 |
| 24051 | 0.0479 | 158983 | 0.0686 | 0.0573 | 4 | 110 | 0.0363 | 0.2 |
| 190640 | -0.0402 | 143785 | -0.0845 | 0.0583 | 3 | 178 | 0.0168 | 0.119 |
| 10607 | 0.0999 | 2984 | 0.0348 | 0.0589 | 1 | 25.8 | 0.0388 | 0.163 |
| 24051 | 0.0479 | 2970 | 0.0767 | 0.0606 | 2 | 233 | 0.00858 | 0.112 |
| 141681 | 0.0895 | 3033 | 0.0443 | 0.063 | 2 | 44.7 | 0.0447 | 0.225 |
| 141681 | 0.0895 | 3733 | 0.0445 | 0.0631 | 4 | 410 | 0.00976 | 0.23 |
| 141681 | 0.0895 | 92 | 0.048 | 0.0655 | 2 | 131 | 0.0153 | 0.195 |
| 140120 | 0.142 | 144403 | 0.0305 | 0.0658 | 1 | 48 | 0.0209 | 0.101 |
| 24051 | 0.0479 | 25998 | 0.0965 | 0.068 | 1 | 251 | 0.00398 | 0.0703 |
| 24051 | 0.0479 | 176155 | 0.0978 | 0.0685 | 1 | 130 | 0.00771 | 0.172 |
| 24051 | 0.0479 | 158756 | 0.103 | 0.0701 | 1 | 79.4 | 0.0126 | 0.206 |
| 140120 | 0.142 | 2984 | 0.0348 | 0.0702 | 1 | 174 | 0.00576 | 0.057 |
| 141681 | 0.0895 | 198836 | 0.0561 | 0.0709 | 1 | 156 | 0.0064 | 0.0664 |
| 5141 | 0.12 | 144366 | 0.0443 | 0.0728 | 1 | 24 | 0.0416 | 0.193 |
| 141681 | 0.0895 | 5150 | 0.0603 | 0.0734 | 1 | 180 | 0.00555 | 0.0482 |
| 5141 | 0.12 | 144425 | 0.0466 | 0.0747 | 2 | 28.3 | 0.0706 | 0.11 |
| 141681 | 0.0895 | 159960 | 0.0651 | 0.0763 | 1 | 64.1 | 0.0156 | 0.145 |
| 10607 | 0.0999 | 150951 | 0.0603 | 0.0776 | 2 | 110 | 0.0182 | 0.0903 |
| 5141 | 0.12 | 198836 | 0.0561 | 0.0819 | 1 | 81.1 | 0.0123 | 0.0892 |
| 190640 | -0.0402 | 177789 | -0.192 | 0.0879 | 2 | 102 | 0.0197 | 0.186 |
| 10607 | 0.0999 | 172141 | 0.0775 | 0.088 | 1 | 98.2 | 0.0102 | 0.112 |
| 24051 | 0.0479 | 4240 | 0.175 | 0.0915 | 4 | 177 | 0.0226 | 0.0719 |
| 140120 | 0.142 | 150951 | 0.0603 | 0.0924 | 1 | 33.7 | 0.0297 | 0.178 |
| 24051 | 0.0479 | 196475 | 0.183 | 0.0935 | 1 | 78.1 | 0.0128 | 0.143 |
| 140120 | 0.142 | 179145 | 0.0639 | 0.0952 | 1 | 119 | 0.00843 | 0.368 |
| 5141 | 0.12 | 172141 | 0.0775 | 0.0963 | 3 | 140 | 0.0214 | 0.103 |
| 5141 | 0.12 | 7293 | 0.086 | 0.101 | 1 | 53.5 | 0.0187 | 0.269 |
| 140120 | 0.142 | 172072 | 0.076 | 0.104 | 1 | 63.7 | 0.0157 | 0.121 |
| 24051 | 0.0479 | 58 | 0.227 | 0.104 | 1 | 225 | 0.00444 | 0.0677 |
| 5141 | 0.12 | 158756 | 0.103 | 0.111 | 1 | 38.4 | 0.026 | 0.131 |
| 140120 | 0.142 | 25998 | 0.0965 | 0.117 | 1 | 14.5 | 0.0692 | 0.106 |
| 24051 | 0.0479 | 190204 | 0.287 | 0.117 | 1 | 192 | 0.00521 | 0.073 |
| 140120 | 0.142 | 4318 | 0.113 | 0.127 | 1 | 51.6 | 0.0194 | 0.0885 |
| 141681 | 0.0895 | 58 | 0.227 | 0.143 | 3 | 330 | 0.00908 | 0.146 |
| 141681 | 0.0895 | 143896 | 0.263 | 0.153 | 6 | 289 | 0.0208 | 0.237 |

| | | | | | | | | |
|---|---|---|---|---|---|---|---|---|
| 140120 | 0.142 | 22901 | 0.167 | 0.154 | 1 | 141 | 0.00707 | 0.231 |
| 141681 | 0.0895 | 190204 | 0.287 | 0.16 | 1 | 109 | 0.00919 | 0.139 |
| 10607 | 0.0999 | 143896 | 0.263 | 0.162 | 1 | 152 | 0.00656 | 0.138 |
| 141681 | 0.0895 | 153600 | 0.3 | 0.164 | 1 | 56.7 | 0.0176 | 0.169 |
| 5141 | 0.12 | 58 | 0.227 | 0.165 | 1 | 27 | 0.0371 | 0.0534 |
| 26266 | -0.538 | 175466 | -0.052 | 0.167 | 1 | 117 | 0.00857 | 0.249 |
| 10607 | 0.0999 | 175650 | 0.301 | 0.173 | 1 | 64.2 | 0.0156 | 0.0415 |
| 140120 | 0.142 | 58 | 0.227 | 0.179 | 1 | 42.7 | 0.0234 | 0.159 |
| 26266 | -0.538 | 7287 | -0.0632 | 0.184 | 3 | 137 | 0.0219 | 0.171 |
| 26266 | -0.538 | 142171 | -0.0985 | 0.23 | 2 | 58 | 0.0345 | 0.214 |
| 26266 | -0.538 | 190907 | -0.107 | 0.24 | 1 | 90 | 0.0111 | 0.102 |

### Supplementary Figures

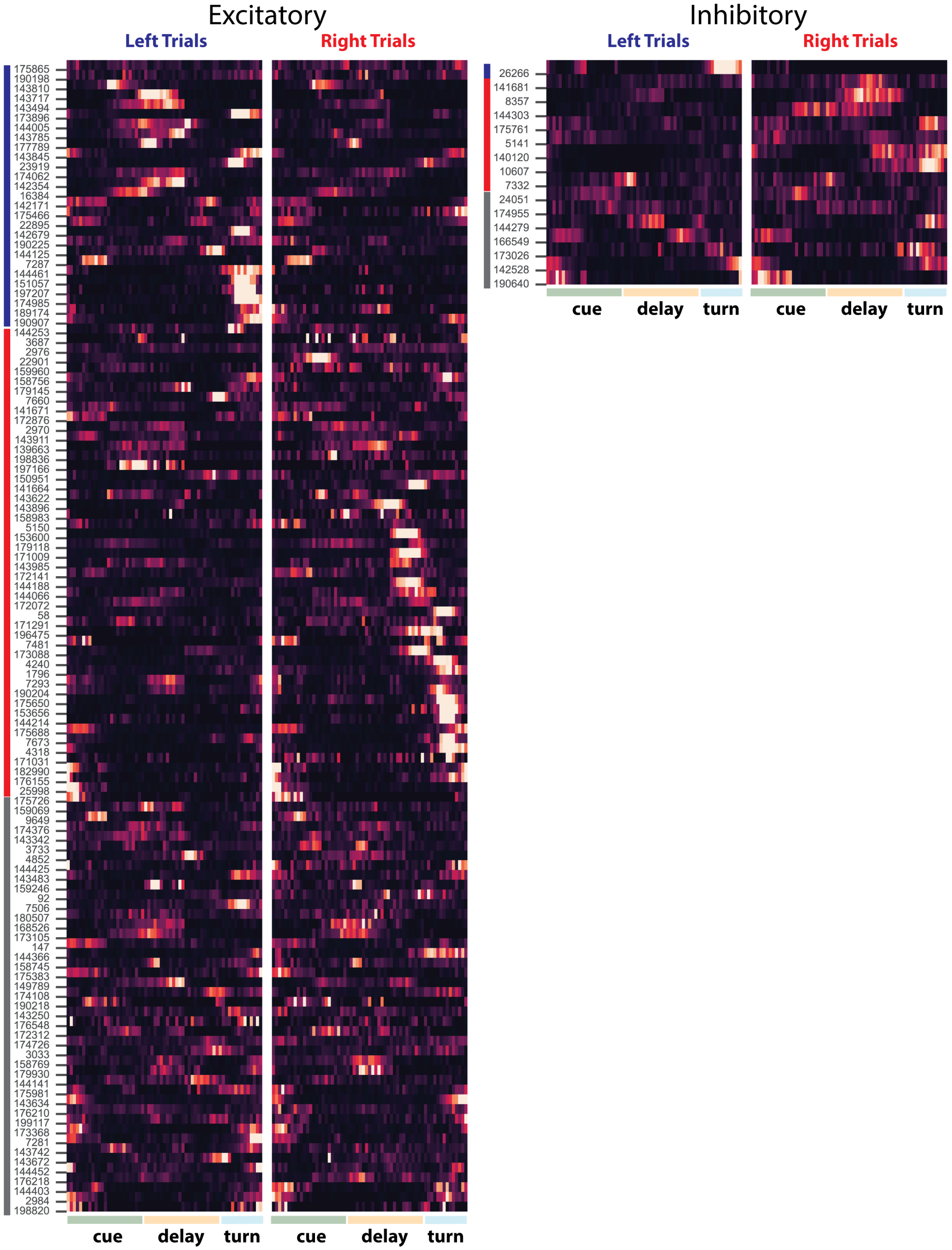


**Supplementary Fig. 1: Trial average neuron activity.** Each row shows trial-average activity of a functionally-characterized neuron. Row labels are neuron IDs corresponding to the CATMAID EM database. Left and right columns show activity on left and right trials, respectively. Activity was averaged over the last 4 behavioral sessions, then normalized by the mean activity rate over both trial types for each neuron separately. Neurons were sorted first by selectivity (left, right, non-selective, indicated by colored bars on y-axis) then by timepoint of maximum mutual information with trial type.

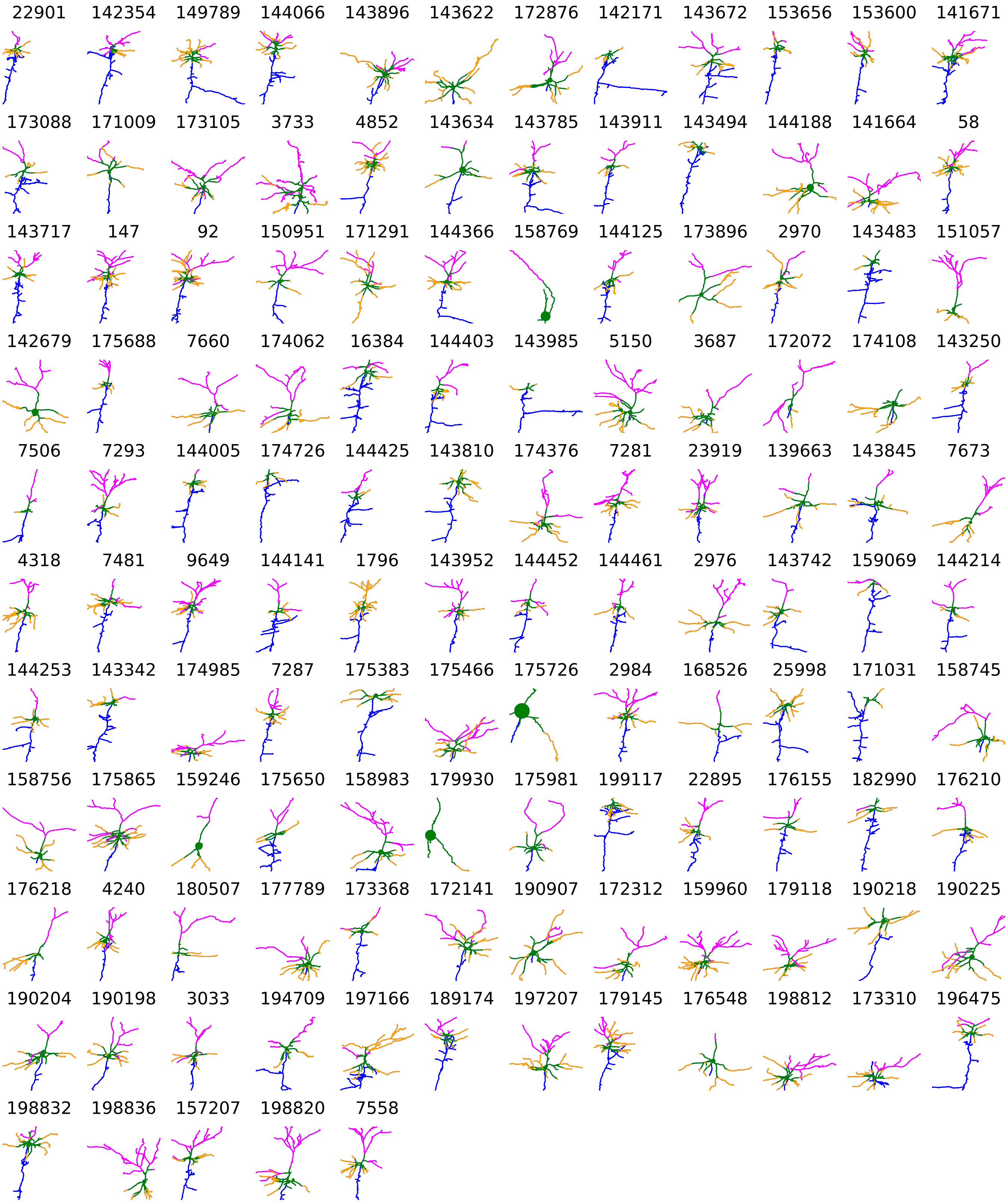


**Supplementary Fig. 2: Excitatory Neuron Morphologies.** Traced neuron morphologies are shown for functionally-characterized excitatory pyramidal neurons in PPC. Axon – blue, proximal dendrites – green, apical dendrites – magenta, basal dendrites – orange. Labels are neuron IDs corresponding to CATMAID database.

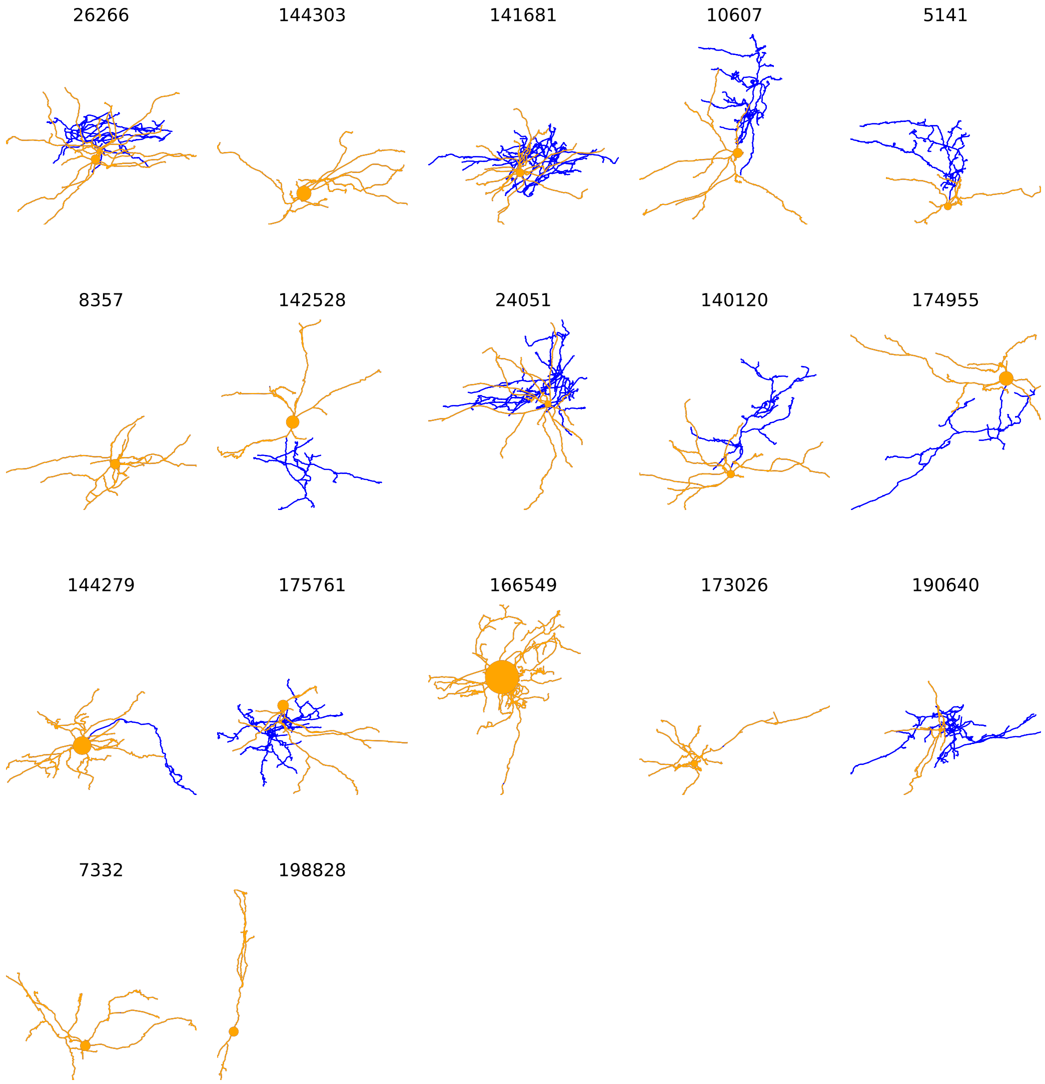


**Supplementary Fig. 3: Inhibitory Neuron Morphologies.** Traced neuron morphologies are shown for functionally-characterized inhibitory neurons in PPC. Axon – blue, dendrites – orange. Labels are neuron IDs corresponding to CATMAID database.

**a** **Decoding accuracy**

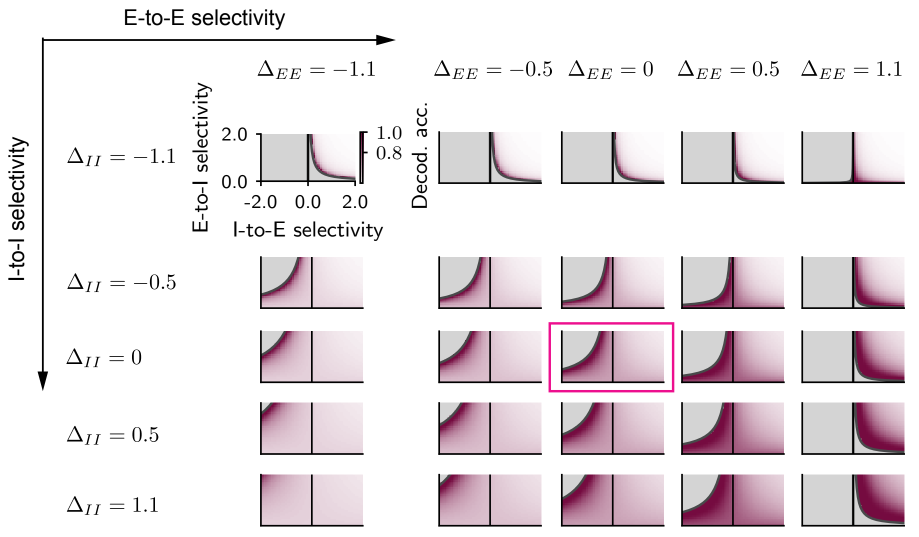


**b** **Relative decoding accuracy**

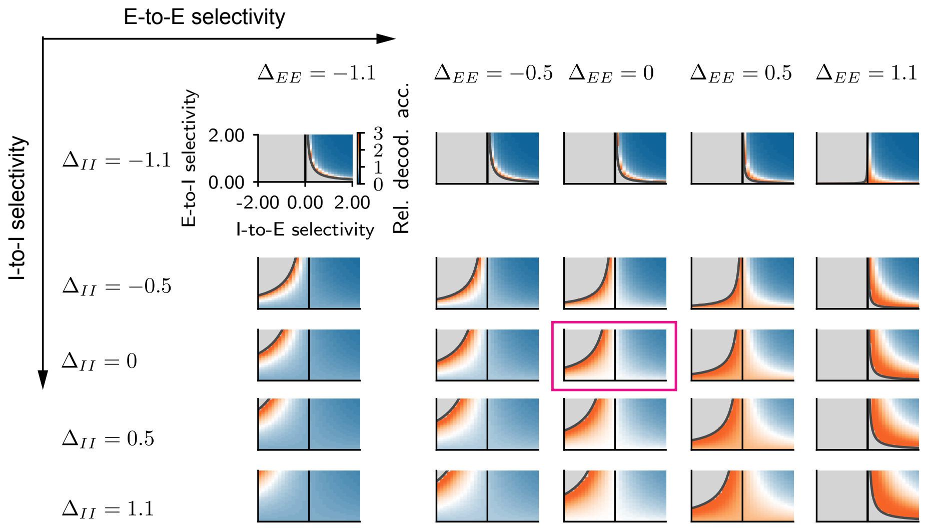


**Supplementary Fig. 4: Dependence of the trial-type decoding accuracy on E-to-E and I-to-I selectivity in the linear rate model. (a)** Dependence of the absolute trial-type decoding accuracy as a function of the selectivity of the four connection types. Each subplot shows the dependence on E-to-I and I-to-E selectivity; different subplots correspond to different values of the E-to-E and I-to-I connection selectivity. The subplot highlighted by a red box corresponds to the one showed in Ext. Data Fig. 4d. In the linear rate model (Fig 4a-d), the region corresponding to selective E-to-I and anti-selective I-to-E (left quadrant) becomes linearly unstable when the E-to-E or I-to-I selectivities satisfy $(1-\Delta_{EE})(1+\Delta_{II}) < 0$. **(b)** Same as **(a)** for the relative decoding accuracy (see Fig. 4c,d).

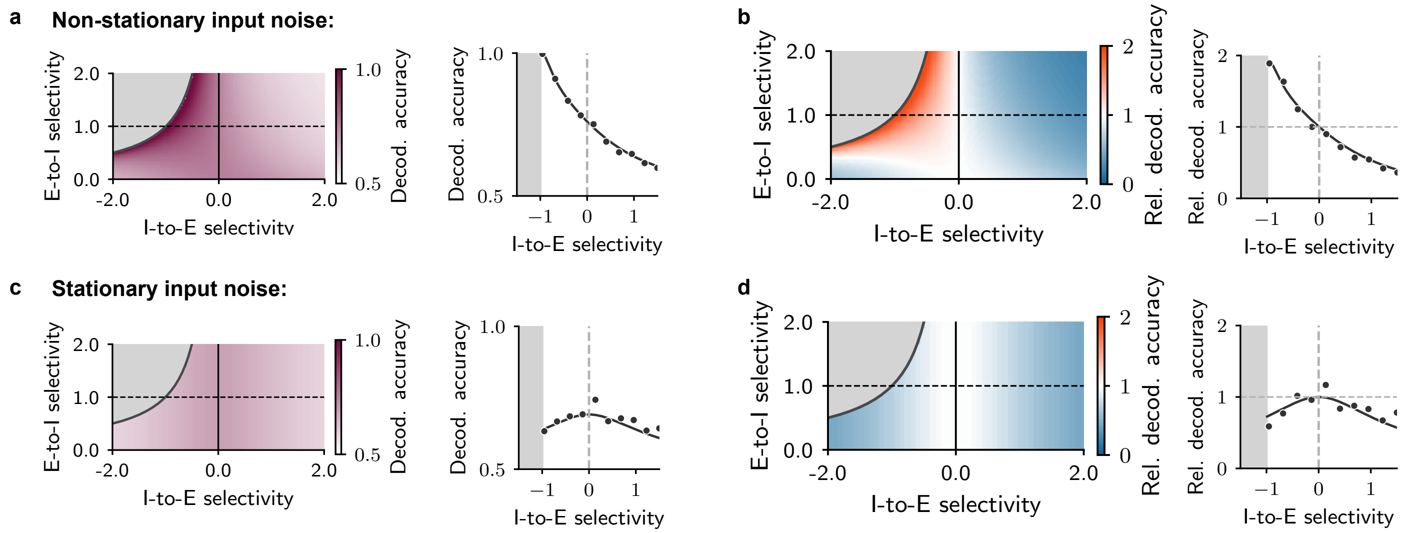


**Supplementary Fig. 5: Trial-type decoding accuracy in presence of input noise in the linear rate model. (a)-(b)** Absolute and relative trial-type decoding accuracy for non-stationary input noise, i.e. input noise that varies in time. The noise affects only the excitatory neurons, which are the neurons that receive the external trial-specific input. **(b)-(c)** Same as **(a)-(b)** for stationary input noise, i.e. noise that affects the external input on the excitatory neurons on a single-trial basis, but is otherwise constant in time.
